# Respiratory complex and tissue lineage drive mutational patterns in the tumor mitochondrial genome

**DOI:** 10.1101/2020.08.18.256362

**Authors:** Alexander N. Gorelick, Minsoo Kim, Walid K. Chatila, Konnor La, A. Ari Hakimi, Barry S. Taylor, Payam A. Gammage, Ed Reznik

## Abstract

Mitochondrial DNA (mtDNA) encodes essential protein subunits and translational machinery for four distinct complexes of oxidative phosphorylation (OXPHOS). Using repurposed whole-exome sequencing data, we demonstrate that pathogenic mtDNA mutations arise in tumors at a rate comparable to the most common cancer driver genes. We identify OXPHOS complexes as critical determinants shaping somatic mtDNA mutation patterns across tumor lineages. Loss-of-function mutations accumulate at an elevated rate specifically in Complex I, and often arise at specific homopolymeric hotspots. In contrast, Complex V is depleted of all non-synonymous mutations, suggesting that mutations directly impacting ATP synthesis are under negative selection. Both common truncating mutations and rarer missense alleles are associated with a pan-lineage transcriptional program, even in cancer types where mtDNA mutations are comparatively rare. Pathogenic mutations of mtDNA are associated with substantial increases in overall survival of colorectal adenocarcinoma patients, demonstrating a clear functional relationship between genotype and phenotype. The mitochondrial genome is therefore frequently and functionally disrupted across many cancers, with significant implications for patient stratification, prognosis and therapeutic development.

## Introduction

Somatic mutations are the underlying drivers of malignancy in cancer, and the identification and characterization of recurrent, functional somatic events has been the capstone goal of cancer genomics. Genomic searches for recurrent driver mutations have focused on the nuclear exome or subsets thereof, motivated by the observation that recurrent mutations are concentrated in the coding regions of a subset of nuclear-DNA-encoded genes. This targeted approach has powered the discovery of common and rare driver mutations in exonic regions, but by corollary has also left underexplored the overwhelming majority of the genome and the driver events it may harbor. Numerous examples now exist of the prevalence and function of oncogenic mutations beyond the nuclear exome, including mutations to the *TERT* promoter, non-coding RNAs including ribosomal RNA and snRNAs, and enhancers ^1^. A fundamental challenge is therefore to discover new functional somatic alterations beyond the exome with a fixed and limited sequencing capacity.

Somatic mutations in tumors commonly target human mitochondrial DNA (mtDNA)^2–6^, affecting both the thirteen essential protein components of four distinct complexes (CI, CIII, CIV, and CV) in oxidative phosphorylation (OXPHOS) as well as the non-coding RNA (22 tRNAs, 2 rRNAs) necessary for mtDNA translation. (**Fig. 1a**). Despite abundant pharmacological, genetic, and clinical data demonstrating that perturbation of different OXPHOS complexes (referred to in shorthand as complexes) produce distinct cellular adaptations ^7,8^, the importance of each complex in shaping mtDNA mutation patterns in cancer is unknown. Because mtDNA is not commonly targeted by exome sequencing panels, prior analyses of mtDNA mutations have relied on cohorts profiled with whole genome sequencing, with consequently diminished statistical power to detect recurrent patterns of mutations relative to exome sequencing studies^8^. However, due to the extremely high copy number and off-target hybridization rate of mtDNA, mtDNA reads are abundant in widely-available exome sequencing of tumors^9^. Mitochondrial DNA therefore represents an opportunity for discovery through repurposing of existing exome sequencing data.

**Fig. 1:**
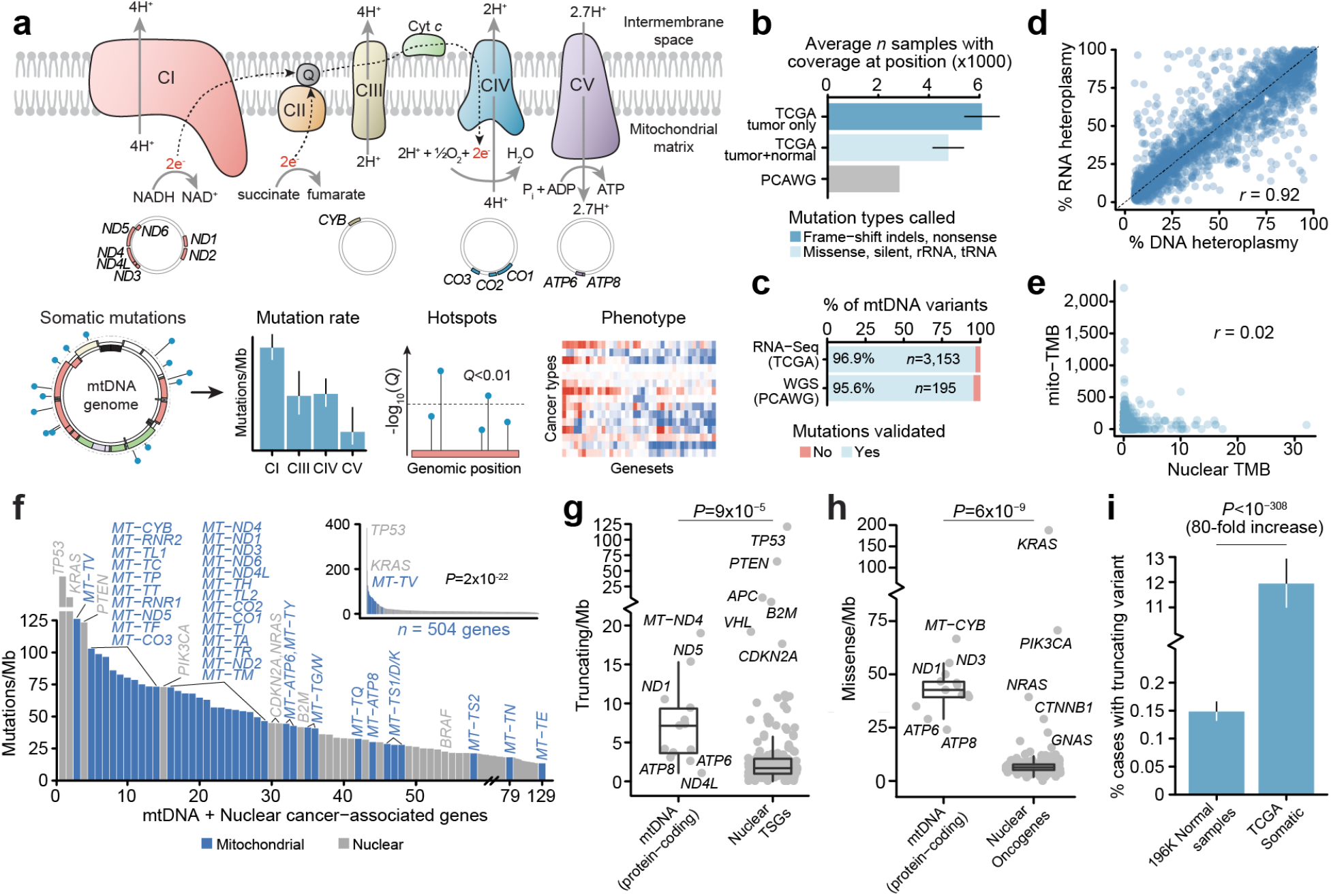
mtDNA mutations are among the most frequent genomic alterations in cancer. **a)** Schematic of oxidative phosphorylation (OXPHOS) system and project workflow. Top row, complexes I-V and their reactions. Center row: mtDNA genomic regions encoding protein subunits of the associated OXPHOS complex. Bottom row, overview of project workflow, in which somatic mutations in mtDNA genes are used to explore inter-complex differences, mutational recurrence and transcriptional phenotype associated with mitochondrial dysfunction. **b)** Average number of tumors with sufficient coverage to call variants at a mtDNA position. Truncating mutations were assumed to be somatic and therefore allowed for tumor-only variant-calling (dark blue), whereas non-truncating (protein-coding non-truncating, tRNA and rRNA mutations) required sufficient coverage in both tumor and matched normal samples (light blue). Gray, the number of whole-genome sequenced (WGS) samples from PCAWG for comparison. **c)** The percentages of variants called from off-target reads which were validated in either RNA-Seq or WGS data from the same tumors. **d)** The correlation between variant heteroplasmy as observed in RNA and DNA-sequencing (*n*=2,575 mutations with coverage ≥30 reads in both DNA and RNA). **e)** The correlation between tumor mutation burden (TMB, Mutations/Mb) among mtDNA (Y-axis) and nuclear-encoded cancer-associated genes (referred to simply as cancer genes) (X-axis), *n*=3,624 well-covered pan-cancer tumors. **f)** Mutation rates (Mutations/Mb) of individual mtDNA-encoded genes (blue) and nuclear-encoded cancer-associated genes (gray). Inset plot: mutation rates among 504 genes with mtDNA genes highlighted. Outer plot: closeup of the inset plot in the region containing all 37 mtDNA genes; commonly-mutated nuclear cancer genes in this region are labeled for reference. **g)** Comparison of truncating mutation rates (truncating variants/Mb) between 13 mtDNA-encoded protein-coding genes and 185 nuclear-encoded TSGs. **h)** Comparison of non-truncating mutation rate (nonsynonymous, non-truncating variants/Mb) between 13 mtDNA protein-coding genes and 168 nuclear oncogenes. **i)** Percentage of patients with truncating mtDNA variants either somatically (in TCGA tumor samples) or germline (among ~200K normal samples).

Here, by utilizing existing exome sequencing data to more than double statistical power of prior analyses, we report that OXPHOS complex, in combination with tissue lineage and mutational consequence, is a critical determinant of mtDNA mutation patterns in cancer. We find that NADH:ubiquinone oxidoreductase (complex I, CI) mutations are strongly enriched for highly pathogenic mutations in specific tissue lineages, whereas ATP synthase (complex V, CV) is broadly depleted of all non-synonymous mutations. We further identify six highly recurrent mtDNA mutation hotspots at specific homopolymer sequence contexts, which collectively account for over 40% of all truncating mutations to mtDNA, as well as recurrent mutations in both protein-coding genes and non-coding RNA elements. These mutations produce a defined, lineage-agnostic transcriptional program and, in specific tumor lineages, associate with both underlying molecular subtypes and clinical outcomes. Our results argue that specific components of mitochondrial respiration are broadly perturbed across many tissue lineages, and that re-analysis of existing genomic data can yield new discoveries in underexplored genomic terrain.

## Results

### mtDNA Mutations in Tumors from Off-target Reads

To study patterns of mtDNA mutations in tumors, we reasoned that the sheer amount of off-target reads aligning to mtDNA in whole-exome sequencing data would be sufficient to call somatic mtDNA mutations in a large proportion of samples. We therefore assembled a dataset of pan-cancer paired tumor and matched-normal exome sequencing samples from the TCGA, *n=*10,132 (**Supplementary Fig. 1a**). Inconsistent sequencing coverage between samples is an inherent limitation to this approach, as variants located in regions without adequate sequencing coverage are not identifiable, and we therefore developed our methodology to be cognizant of the sequencing coverage at each position in each sample (see **Methods**). We focused our analysis on regions of mtDNA in protein-coding genes and genes coding for mitochondrial ribosomal RNAs (rRNAs) and transfer RNAs (tRNAs), excluding the control region (pos 1-576, 16,024-16,569), several known hypervariable loci (pos 302-314; 514-524; 3,106-3,109), and 89 remaining positions not within any genic region from all further analyses (excluded positions listed in **SI table 1**). The combination of an increase in sample size (in TCGA, relative to whole genome cohorts) and high off-target read coverage effectively doubled the number of tumor-associated mtDNA genomes sequenced compared to the largest published dataset of whole-genome sequenced tumor mitochondrial genomes ^3^: On average 6,100 tumors were sequenced at sufficient depth to call mutations at each mtDNA position (mean+/−SD: 5,399-6,800 samples covered at a given position, **Fig. 1b**, **Supplementary Fig. 1b**), compared to 2,836 whole-genome tumor sequences from the PCAWG dataset. When further restricted to regions sequenced at sufficient depth in both tumor and matched-normal samples, each position was covered in 4,769 tumor/normal pairs on average (mean+/−SD: 4,148-5,390 samples).

We implemented a conservative variant calling approach modeled after state-of-the-art methodologies for exome sequencing, in which we took the intersection of two variant callers (MuTect2 ^10^ and an in-house variant caller based on the SAMtools mpileup utility ^11^, see **Methods**). Consistent with prior work, mtDNA variants exhibited a strand-specific enrichment for C>T mutations on the heavy strand and T>C mutations on the light strand (**Supplementary Fig. 1c**). Based on 789 tumor samples from TCGA with whole genome sequences in the PCAWG cohort ^3^, 95.6% of mutation calls from whole exome sequencing validated against published mutation calls from the PCAWG data (**Fig. 1c**). We also evaluated the possibility that nuclear-encoded mitochondrial pseudogenes (NUMTs) could corrupt variant calling. Although both mtDNA and NUMTs are not targeted by exome sequencing, mtDNA is unique in that it exists at orders of magnitude higher copy number in each cell and, critically, is expressed at extremely high levels, whereas NUMTs do not show evidence of significant transcription ^12^. We therefore determined the fraction of somatic mtDNA variants from exome sequencing which could be recapitulated in matched RNA sequencing from the same sample, revealing that 96.9% of such variants were validated. Finally, we observed a strong correlation between DNA and RNA heteroplasmy overall (Pearson’s r = 0.918) (**Fig. 1d**), confirming that the vast majority of observed mutations are expressed and providing further evidence that the mutations called by our approach are not attributable to NUMTs. In total, we identified 4,381 mtDNA mutations from 10,132 tumor samples which were either protein-truncating (*i.e.* frame-shift indels or nonsense mutations); or non-truncating variants (missense, in-frame indels, translation start-site, non-stop, or mutations to tRNA/rRNAs) which were detected in tumor and absent from matched-normal samples. Among a subset of 3,264 paired tumor/normal samples with sufficient coverage to call mtDNA mutations in at least 90% of the mitochondrial genome (32% of tumor/normal pairs in our dataset overall, referred to throughout as “well-covered” samples), 57% (95%CI 56-59%) had at least one mtDNA variant, in agreement with previous estimates for mtDNA mutation incidence in pan-cancer sequencing data ^2^. Consistent with independent mutagenic processes operating in the nuclear and mitochondrial genomes, we observed no correlation between nuclear and mitochondrial mutation burdens pan-cancer or within individual cancer types (**Fig. 1e**, **Supplementary Fig. 1d)**. Furthermore, in colorectal and stomach cancers where microsatellite instability (MSI) is common, the presence of MSI affected mutation burden in the nuclear but not in the mitochondrial genome (**Supplementary Fig. 1e**).

The mutation rate in the coding region of mtDNA is roughly 67.8 mut/Mb, roughly 6-fold higher than the rate in 468 cancer-associated genes in the MSK-IMPACT panel ^13^ of 11.3 mut/Mb (*P* < 10^−308^ (computational limit of detection), two-sided Poisson test). When calculated for each gene, we also observed mtDNA-encoded genes to have enriched mutation rates compared to nuclear-DNA-encoded MSK-IMPACT genes (*P*=2×10^−22^, two-sided Wilcoxon rank sum test): only 2 MSK-IMPACT genes (*TP53, KRAS*) exhibited rates higher than that of the most mutated mtDNA-encoded genes (**Fig. 1f**). Furthermore, the 13 protein-coding mtDNA genes exhibited a 4.2-fold higher rate of truncating variants which disrupt the reading frame (*i.e.* nonsense mutations and frameshift indels) compared to truncating mutations among 185 known tumor suppressor genes in the MSK-IMPACT cohort (which also included splice-site variants which cannot arise in the mitochondrial genome due to the lack of introns) (*P*=9×10^−5^, two-sided Wilcoxon rank sum test) (**Fig. 1g**), and a 6.7-fold higher rate of non-truncating, non-synonymous mutations (collectively referred to here as “missense” mutations) than 168 MSK-IMPACT oncogenes (*P*=6×10^−9^) (**Fig. 1h**). In total, 11.9% of tumors across all cancers (95% CI: 11.0-12.9%) harbored a truncating mtDNA variant absent in the patient’s matched normal sample. In contrast, only 0.15% of normal blood samples exhibited a truncating variant (95% CI: 0.13-0.17%) based on a recent analysis of ~200,000 mtDNA genomes ^14^ (**Fig. 1i)**. The rate of truncating mutations in mtDNA genes in tumors therefore represents an 80-fold increase compared to truncating mutations observed in normal human genomes (**SI table 2.)** Of the 619 truncating mutations we observed, 196 (32%, 95% binomial CI: 28-35%) had >80% heteroplasmy despite underlying infiltration of the bulk tumor by normal stromal and immune cells, indicating that a significant number of tumors are dominated by a highly dysfunctional mitochondrial genotype. Furthermore, high heteroplasmy truncating variants were significantly more common than high-heteroplasmy silent mutations (139/555, 25%, 95% CI: 21-29%) (*P*=0.01, two-sided Fisher’s exact test) under predominantly neutral selection.

### Truncating Mutations Preferentially Target Complex I at Homopolymeric Hotspots

The physiologic response to genetic and pharmacologic inhibition of mitochondrial respiration depends strongly on which mtDNA-encoded complex (CI, CIII, CIV, CV) is disrupted, implicating OXPHOS complex as a potential determinant of selective pressure for mutation. We therefore investigated the somatic mutation rate according to the OXPHOS complex, controlling for the relative length of mtDNA coding for genes in each complex and uneven coverage within each sequenced sample. This revealed a striking dichotomy in the relative enrichment of mutations in each complex. Truncating variants (nonsense mutations and frame-disrupting indels) arose at a 2-fold or greater rate in complex I relative to the other complexes (*P*=0.001 for least significant comparison, two-sided Poisson test) (**Fig. 2a)**. No difference in mutation rate between complexes was observed for silent mutations, consistent with a lack of differential selective pressure for synonymous protein changes (*P*=0.5 for most-significant comparison). Unlike variants in other complexes, truncating variants in CI demonstrated higher heteroplasmy (variant allele frequency) than silent variants (*P*=1×10^−6^, CI; most significant for other complexes, *P*=0.4, two-sided Wilcoxon rank sum test) (**Fig. 2c**). Finally, complex V genes (*MT-ATP6* and *MT-ATP8*) demonstrated significantly lower rates of truncating but not synonymous mutations. The findings above were recapitulated in an independent cohort, composed of a distinct mixture of cancer types, of N=1,951 whole-genome sequenced tumors from the PCAWG dataset, after excluding samples overlapping with our own cohort (**Fig. 2b**). Tumors of different lineages exhibited wide variability in the incidence of truncating mutations, with ≤5% of some cancer types affected by truncating mutations (sarcomas, gliomas), to 20% or greater of other cancer types (renal cell, colorectal, thyroid) (**Fig. 2d**). In renal, thyroid, and colorectal cancers, the high burden of truncating variants was driven by a specific enrichment for mutations to complex I (*Q*-value < 0.01, two-sided McNemar’s test) (**Fig. 2e)**. Truncating variants in these three cancers affected between 20-30% of all samples, corresponding to a prevalence on the same order or exceeding that of common tumor suppressors in these diseases. Taken together, these data indicate that the functional consequence of mtDNA variants and the complex they target are key determinants of the pattern of somatic mtDNA mutations. Additionally, they suggest that disruption of complex V, which would fundamentally impair mitochondrial ATP production independent of the activity of all other OXPHOS complexes, is not tolerated.

**Fig. 2:**
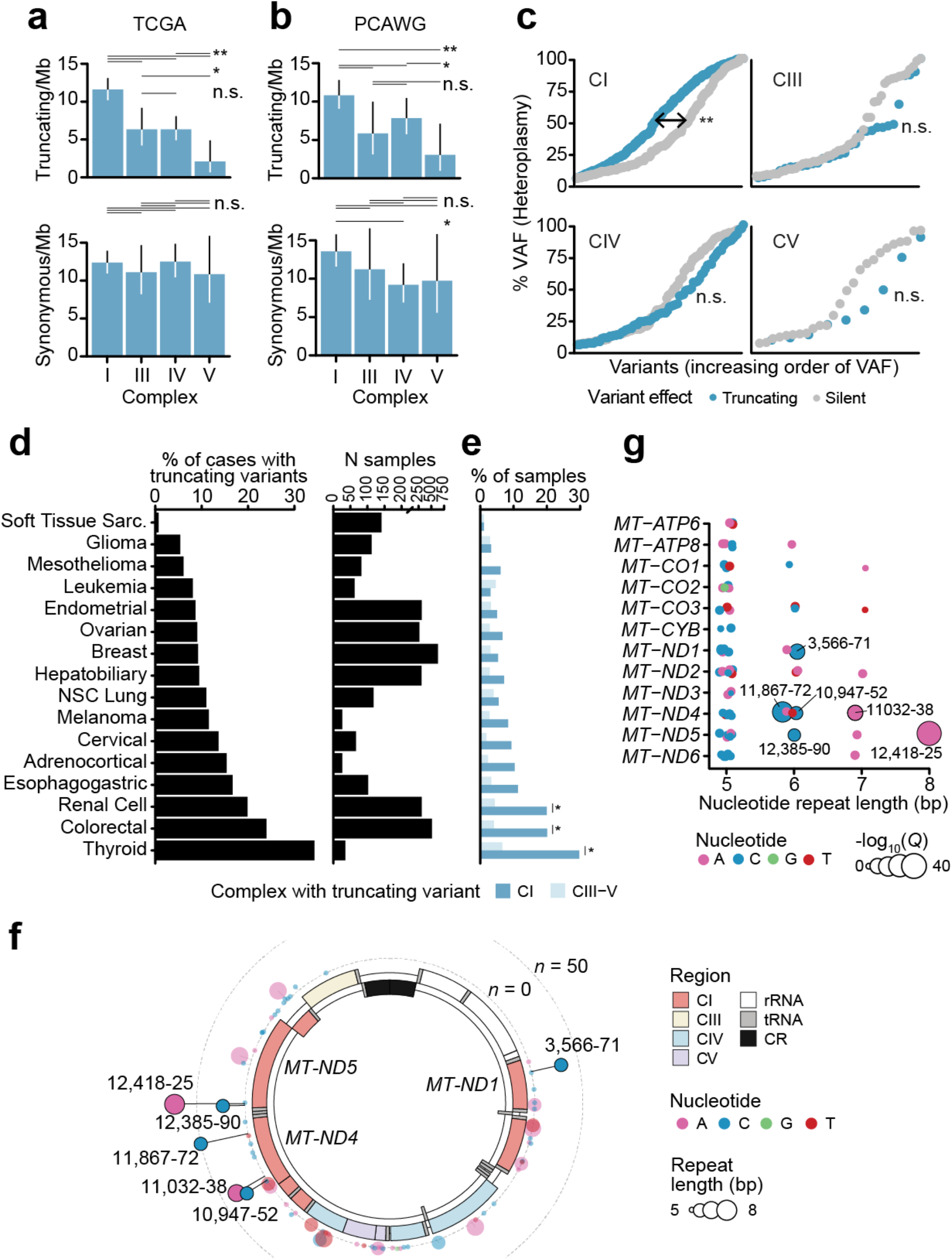
Truncating variants preferentially target complex I. **a)** Comparison of truncating mutation rate (truncating variants/Mb) between OXPHOS complexes I, III, IV, V. Synonymous mutation rates shown below for comparison. Truncating mutations *n*=352; synonymous *n*=475. *P*-values from two-sided Poisson-exact. Single asterisk denotes *P*<0.1; double asterisk *P*<0.01; n.s., not significant. **b)** Validation of analysis in a) using data from n=1,951 whole-genome sequenced tumors from ICGC/PCAWG after removing samples also in TCGA. Truncating mutations *n*=198; synonymous *n*=263. *P*-values and asterisks as in a). **c)** Distributions of truncating and silent mutation heteroplasmy (estimated by variant allele frequency) among variants in OXPHOS complex I, III, IV, or V. Difference in heteroplasmy between truncating and silent mutations calculated by two-sided Wilcoxon rank sum test. CI, *P*=1×10^−6^, not significant for other complexes. **d)** Percentage of tumors with truncating mtDNA variants per cancer type, among well-covered samples. Right, number of well-covered samples per cancer type. **e)** Percentage of samples per cancer type with truncating variants affecting OXPHOS complex I or III-V. Asterisk indicates cancertypes with enriched truncating variants targeting CI compared to CIII-V, *Q*<0.01, two-sided McNemar’s test. **f)** Circular mtDNA genome annotated with 73 homopolymer repeat loci ≥5bp in length. Dot height from the circular mtDNA genome indicates the number of affected samples, dot color indicates the identity of the repeated nucleotide (A, C, G, T), dot width indicates the length of the repeat region (5-8bp). Includes putatively somatic truncating variants with tumor-only sequencing coverage. The 6 solid-color homopolymer loci highlighted were found to be statistically enriched hotspots for frame-shift indels in tumors. **g)** The 73 homopolymer repeat loci arranged by gene and repeat size. Dot width indicates – log_10_(*Q*-value) for enriched frame-shift indels in tumors. The 6 hotspot loci are labeled.

Unexpectedly, we observed that truncating mutations frequently arose at the same genomic locus, analogous to well-described hotspot mutations that accumulate in nuclear cancer driver genes and often reflect selective pressure ^15,16^. These apparently recurrent alleles were exclusively indels, rather than nonsense mutations, characterized by a homopolymeric sequence context. We therefore developed an approach to detecting recurrent mutations at homopolymeric loci by modeling incidence of frame-shift indels at each locus as a function of their base-pair length (see **Methods**). Six single-nucleotide repeat loci (out of seventy three loci of 5 or more base-pairs in length) in *MT-ND1* (c.3,566-3,571, *n =* 32), *MT-ND4* (c.10,947-10,952, *n =* 25; c.11,032-11,038, *n* = 34; and c.11,867-11,872, *n* = 50), and *MT-ND5* (c.12,385-12,390, *n* = 23 and c.12,418-12,425, *n* = 73) accumulated mutations at a rate above null expectation (*Q*-value<0.01, **Fig. 2f**). Homopolymer hotspots only arose at single-nucleotide loci of at least 6 nt in length (*P*=0.0002, two-sided Fisher’s exact test), were composed of A or C homopolymer repeats, and exclusively encoded subunits of complex I. Importantly, other homopolymers of equivalent length (≥6) and nucleotide content exist both in complex I and complex III/IV/V but did not exhibit recurrent mutations, indicating a high degree of specificity to hotspot positions (**Fig. 2g)**. These six homopolymeric repeat loci collectively accounted for 40% of all truncating variants observed in our data (95% binomial CI: 36-44%), and 57% (95% CI: 52-62%) of frame-shift indels overall. Notably, recurrent loss-of-function frameshift indels have been observed at these sites as early driver mutations in rare, often benign renal oncocytomas 17; however we observed mutations at these loci to be a pervasive phenomenon across tumor lineages (**Supplementary Fig. 2a**). Homopolymeric hotspot mutations arose in the PCAWG cohort (after excluding any samples overlapping with our cohort) at a rate highly consistent with the TCGA cohort (Pearson’s *r* = 0.95), indicating that the indels detected in TCGA at hotspot loci were not artifacts due to calling variants in microsatellite regions with poor coverage (**Supplementary Fig. 2b**). Moreover, the three most prevalently mutated of these homopolymer loci in our dataset (c.11,032-11,038, c.11,867-11,872, c.12,418-12,425) intersected with 100-bp-long windows enriched for frameshift indels identified in an analysis of 616 pediatric and 2,202 adult tumors (527 of which were from TCGA), highlighting the power of our approach to resolve focal, recurrent alterations ^18^. Although mutations at homopolymeric tracts have not been widely described in the germline literature, the most recurrent hotspot (*MT-ND5* c.12,418-12,425) has been previously reported as the site of a germline frame-shift deletion (A12425del) in a pediatric patient, where the *de novo* heteroplasmic deletion resulted in mitochondrial myopathy and renal failure^19^.

### Non-synonymous mutations and RNA variants arise as rare recurrent alleles with elevated pathogenicity

The bulk of somatic variants we observed in mtDNA were non-truncating, non-synonymous mutations, including missense mutations, in-frame indels, translation start site mutations and non-stop mutations (collectively variants of unknown significance or VUS, 73.2% of *n*=4,381 variants, **Fig. 3a**). Interestingly, non-synonymous variants were again depleted in CV relative to other complexes, suggesting that CV is intolerant both to truncating variants and to presumably less-disruptive non-synonymous mutations. Using the APOGEE framework to evaluate the functional consequence of mutations to protein-coding mtDNA genes ^20^, we found that somatic VUSs were twice as likely to be predicted pathogenic compared to germline polymorphisms observed among ~200K normal samples from the HelixMTdb dataset (39.5% of somatic-only variants compared to 20.4% of germline-arising, *P*=6×10^−14^, two-sided Wilcoxon rank sum test, **Fig. 3b**). Furthermore, when considering all possible mtDNA variants excluding germline polymorphisms (*i.e.* the complete set of all possible somatic variants), VUSs observed in tumors were more pathogenic than the set of possible somatic variants which never arose in tumors, suggesting that somatic VUSs are more pathogenic than expected by random chance. We next evaluated the tendency for VUSs to target specific complexes of the ETC (this necessarily reduced the types of VUSs to protein-coding variants, including missense, in-frame indels, and a small number of translation start site and nonstop mutations). In contrast to truncating variants, protein-coding VUSs were most frequent in CIII (*P*=1×10^−7^ for least significant comparison, two-sided Poisson test, **Fig. 3c**), whose functional integrity as a site for ubiquinol oxidation has recently been described as essential for tumor cell proliferation^21^, although as with truncating variants VUSs to CV subunits were still depleted compared to the other complexes (*P*=0.01 for least significant comparison). These observations were validated using data from PCAWG (**Fig. 3d**). Together, these findings suggest that tumors preferentially accumulate somatic missense mtDNA mutations in a manner dictated by OXPHOS complex, possibly driven by their capacity to disrupt mitochondrial function due to their elevated pathogenicity. Furthermore, they support the hypothesis that a purifying selection exists against variants (both truncating and VUSs) that compromise physiological function of complex V/ATP Synthase.

**Fig. 3:**
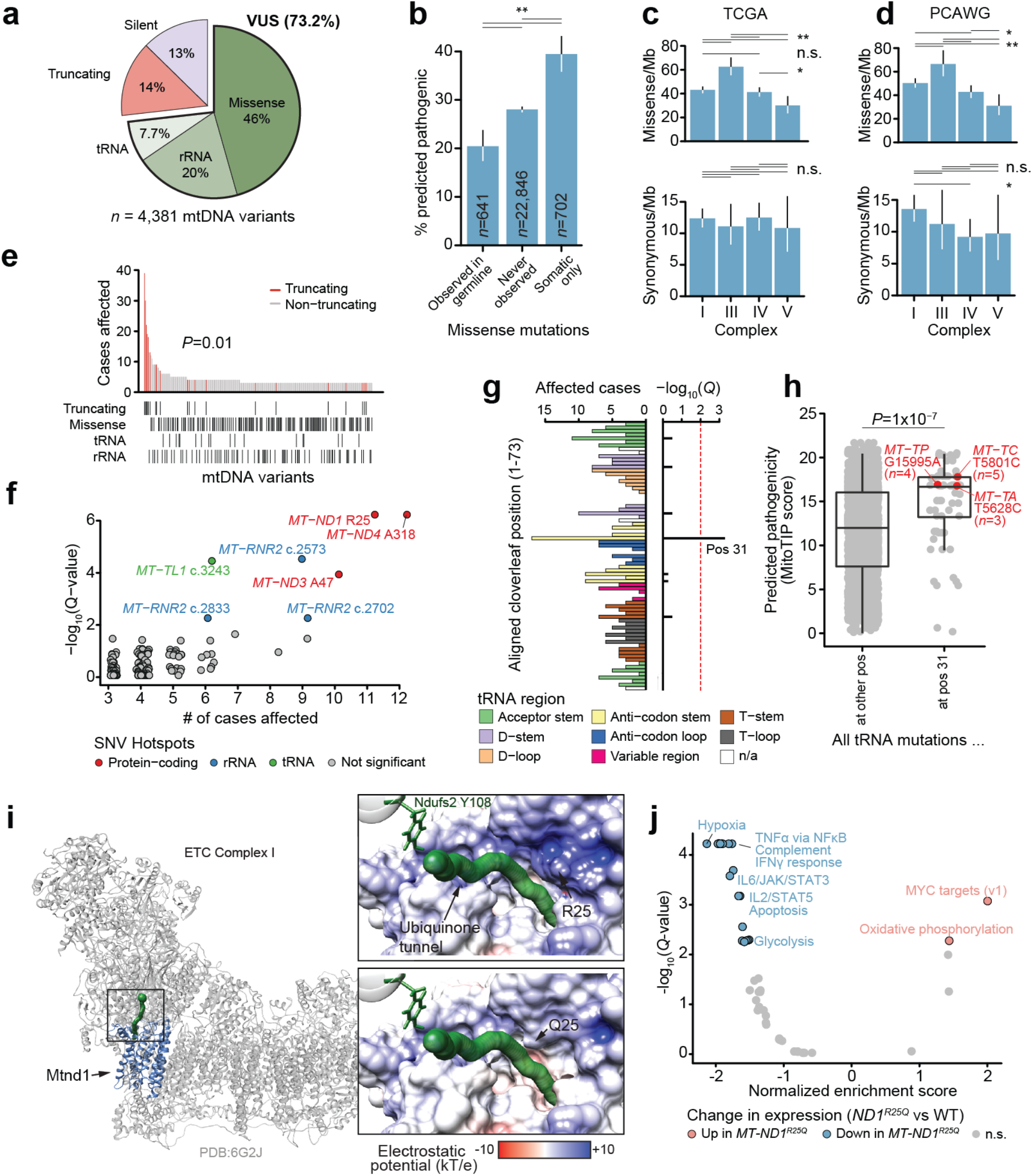
Non-truncating mtDNA mutations arise as rare recurrent alleles in protein-coding and RNA elements. **a)** The proportion of truncating, synonymous and VUS somatic mtDNA mutations in this study. VUSs are further classified into missense protein-coding variants, or mutations to rRNA or tRNA genes. **b)** Comparison of the percentage of unique VUSs predicted to be pathogenic by APOGEE ^20^ between somatic variants which (1) were ever observed to also arise as germline variants among ~200K normal samples from HelixMTdb; (2) were never observed somatically mutated; or (3) were only observed as somatic mutations. All indicated comparisons were statistically significant with P<10^−4^. **c)** Comparison of missense mutation rate (missense variants/Mb) between OXPHOS complexes I, III, IV, V. Synonymous mutation rates shown below for comparison. Missense mutations *n*=1,718; synonymous *n*=475. *P*-values from two-sided Poisson-exact. Single asterisk denotes *P*<0.1; double asterisk *P*<0.01; n.s., not significant. **d)** Validation of analysis in a) using data from n=1,951 whole-genome sequenced tumors from ICGC/PCAWG after removing samples also in TCGA. Missense mutations *n*=1073; synonymous *n*=263. *P*-values and asterisks as in a). **e)** Rare-recurrent alleles are primarily non-truncating variants. Top portion, number of samples with the given mtDNA mutant allele in decreasing order of prevalence (mutations called in samples with adequate tumor and normal coverage). Bottom portion, tracks indicate consequence of corresponding variant in top portion. **f)** Individual base-pair positions in mtDNA with somatic single-nucleotide variants (SNVs) in ≥5 tumors, and their statistical enrichment for mutations. Hotspot positions with *Q*<0.01 are colored by the type of gene in which they arise (protein-coding, rRNA or tRNA). Select hotspots are labeled with their genomic positions (for mutations in tRNAs and rRNAs) or residue (protein coding genes). **g)** Prevalence of SNVs in tRNA genes, aligned to their positions in the folded tRNA cloverleaf structure. Bottom portion, number of samples with SNVs at the given tRNA cloverleaf position across all tRNAs. Top portion, statistical enrichment for the aligned position for mutations. **h)** Mutations at tRNA cloverleaf structural position 31 have greater predicted pathogenicity scores (based on MitoTIP ^39^) compared to all possible mutations at other positions. tRNA mutations at position 31 affecting ≥2 samples are highlighted. *P*-value from two-sided Wilcoxon rank sum test. To reduce image size, a random selection of 5% of the mutations not at position 31 are plotted (*P*-value based on the complete set of mutations). **i)** The Mtnd1 R25Q mutation lies at a critical region of complex I near the entrance to the ubiquinone binding tunnel (dotted green path), likely affecting its capacity for binding ubiquinone. Larger view: The complete mammalian complex I structure (gray) highlighting Mtnd1 (blue), and the ubiquinone binding tunnel (green) and binding site (large green sphere); black box indicates the region in the closeup view. Closeups, the predicted surface electrostatic potential of Mtnd1 (top) wild-type and R25Q mutant (bottom), proximal to the ubiquinone binding tunnel (green), leading to its binding site at Ndufs2 Y108. **j)** Differentially expressed mSigDB Hallmark genesets between colorectal tumors with *MT-ND1* R25Q and those without non-silent somatic mtDNA variants (*i.e.* wild-type). Normalized enrichment score (NES) and adjusted P-values based on gene set enrichment analysis (GSEA) using the fgsea R package ^44^.

Single nucleotide variants (SNVs) were far less recurrent than homopolymer indels (*P*=0.01, two-sided Wilcoxon rank sum test among distinct variants mutated in >=3 tumors, **Fig. 3e**). However, we nevertheless observed a small number of loci with recurrent non-truncating variants. recurrent mutant loci. We developed a statistical test for recurrence of these loci, and identified 7 SNV hotspots in the mitochondrial genome (*Q*<0.01, **Fig. 3f**), including 3 in protein-coding genes (all in complex I), 3 in ribosomal RNAs (all in *MT-RNR2*), and 1 in a tRNA (*MT-TL1*) (**see Methods**). In contrast to the high fraction of truncating mutations which are explained by a relatively small number of hotspot alleles, hotspot SNV mutations collectively accounted for 1.6% of all VUSs; the vast majority of VUSs were non-recurrent, usually arising in a single sample. Furthermore, 0/33 mutations arising at the three protein-coding hotspot positions were nonsense mutations introducing an early stop codon, suggesting either the mutagenic mechanism generating homopolymeric indel hotspots has a high degree of specificity, or that truncating hotspots themselves may engender unique phenotypes beyond conventional loss-of-function.

Mitochondrial tRNAs (mt-tRNAs) are commonly mutated in the context of germline mitochondrial disease. Interestingly, the somatic hotspot *MT-TL1^A3243G^* (somatically mutated in 6 patients) is also the causative variant of around 80% of MELAS disease cases and approximately 30% of all mtDNA disease ^22,23^. We additionally observed mutations clustered in adjacent positions 3242 (*n* = 5) and 3244 (*n* = 4, recently described as a recurrent mutation in Hürthle cell carcinoma of the thyroid ^24^), suggesting that recurrent mutations in *MT-TL1* could affect a common secondary structure element. Mitochondrial tRNAs adopt a relatively conserved cloverleaf structure upon folding, and mutations to mt-tRNAs are known to disrupt the function of specific secondary structure elements. We therefore sought to test whether any positions of the tRNA cloverleaf structure were enriched for somatic mutations in tumors. We aligned all tRNA mutations according to their position in the canonical mitochondrial tRNA structure and developed a statistical approach to identify enrichment in specific secondary structure elements (see **Methods**). This analysis identified position 31 in the anti-codon stem of the folded tRNA molecule as a site of recurrent mutation across mt-tRNAs (*Q*=4.7×10^−4^, **Fig. 3g**), which we further validated using the non-TCGA subset of PCAWG samples (*Q*=0.014, **Supplementary Fig. 3a**). Interestingly, position 31 was observed to be mutated at an 8-fold higher rate in tRNAs encoded on the light-strand (*e.g. MT-TC*, *n*=5; *MT-TP*, *n*=4; *MT-TA*, *n*=3) compared to heavy-strand-encoded tRNAs (*P*=2×10^−4^, two-sided Fisher’s exact test). As a group, mutations at structural position 31 were predicted to be more pathogenic by MITOTIP relative to mutations at other tRNA positions (**Fig. 3h**), and in the case of *MT-TA^T5628C^* (*n*=3) are associated with the mitochondrial disease chronic progressive external ophthalmoplegia (CPEO) ^25^. In analogy to the recurrent mutation of conserved amino acid residues in domains of homologous proteins ^26^ or within 3-dimensional regions of folded protein structures ^27^, these data suggest that specific structural features of mt-tRNAs may undergo recurrent mutation and impair normal mitochondrial physiology.

To understand the potential function of rare protein-coding SNV hotspots in mtDNA, we focused on a recurrent mutation at *MT-ND1*^R25^, which was identified somatically in 11/10,132 TCGA patients (0.11%), and 5/2,836 PCAWG patients (0.18%). All 16 instances resulted in a substitution of arginine (R) with glutamine (Q), encoded by a G>A substitution at position 3380. *MT-ND1*^R25Q^ was previously described in a case report as the causative variant in the development of MELAS in a mitochondrial disease patient ^28^, but was never observed among ~200K normal samples, where the mutant alleles at residue R25 always produced synonymous mutations (A3381G, *n*=57). Residue R25 is conserved across vertebrates ^28^, and is part of a cluster of charged residues in complex I which form a structural bottleneck in the ubiquinone binding tunnel leading to the Q binding site^29^. This led us to hypothesize that the R25Q mutation could potentially disrupt the site, impacting ubiquinone: complex I binding kinetics and/or Q-site substrate specificity, impeding the downstream electron transport chain. We therefore modelled the effect of *MT-ND1*^R25Q^ using a recent, high resolution structure of the mammalian. This analysis highlighted changes to the local charge environment due to loss of the relatively bulky, positively charged arginine sidechain. Due to the location of this substitution within the Q binding tunnel, this is predicted to significantly impact function (**Fig. 3i**). Focusing on colorectal tumors, which demonstrated the largest numbers of tumors harboring *MT-ND1^R25Q^* (*n=*8 tumors total), we examined whether the presence of *MT-ND1^R25Q^* was associated with a particular transcriptional signature. Relative to mtDNA-wild-type tumors, we observed that *MT-ND1^R25Q^* tumors were characterized by upregulation of MYC targets and oxidative phosphorylation, and downregulation of gene signatures associated with hypoxia, IL2/STAT5 signaling, TNFα Signaling via NFκB (**Fig. 3j**). These data suggest that *MT-ND1^R25Q^* promotes a transcriptional phenotype characterized by increased mitochondrial metabolism and suppressed expression of innate immune genes.

### Mitochondrial genotype underlies a lineage-agnostic transcriptional program

Given the lineage specificity underlying both truncating variants and truncating/SNV hotspots, we studied the overall burden of distinct classes of mtDNA variants (*i.e.* producing a truncating, missense, synonymous, tRNA or rRNA variant) across cancer types. Restricting our analysis to well-covered samples including coverage over all homopolymeric hotspots (see **Methods**), we found that the fraction of mutant samples across cancer types ranged from approximately 23% of leukemias (95% binomial CI: 13-35%) to as high as 80% of thyroid cancers (95% CI: 63-92%) (**Fig. 4a**). Moreover, we observed no correlation between the fraction of well-covered samples in a cancer type and the proportion of samples with a somatic mtDNA mutation (**Supplementary Fig. 4a)**, indicating that the highly variable incidence of different somatic variants across cancer types was not biased by their differing sequencing coverages. This extensive variation suggests tumor lineages may be subject to different degrees of selection for or against mtDNA mutations, consistent with the extensive variability of dN/dS ratios previously described in somatic mtDNA mutations derived from whole genome sequencing of the TCGA ^5^.

**Fig. 4:**
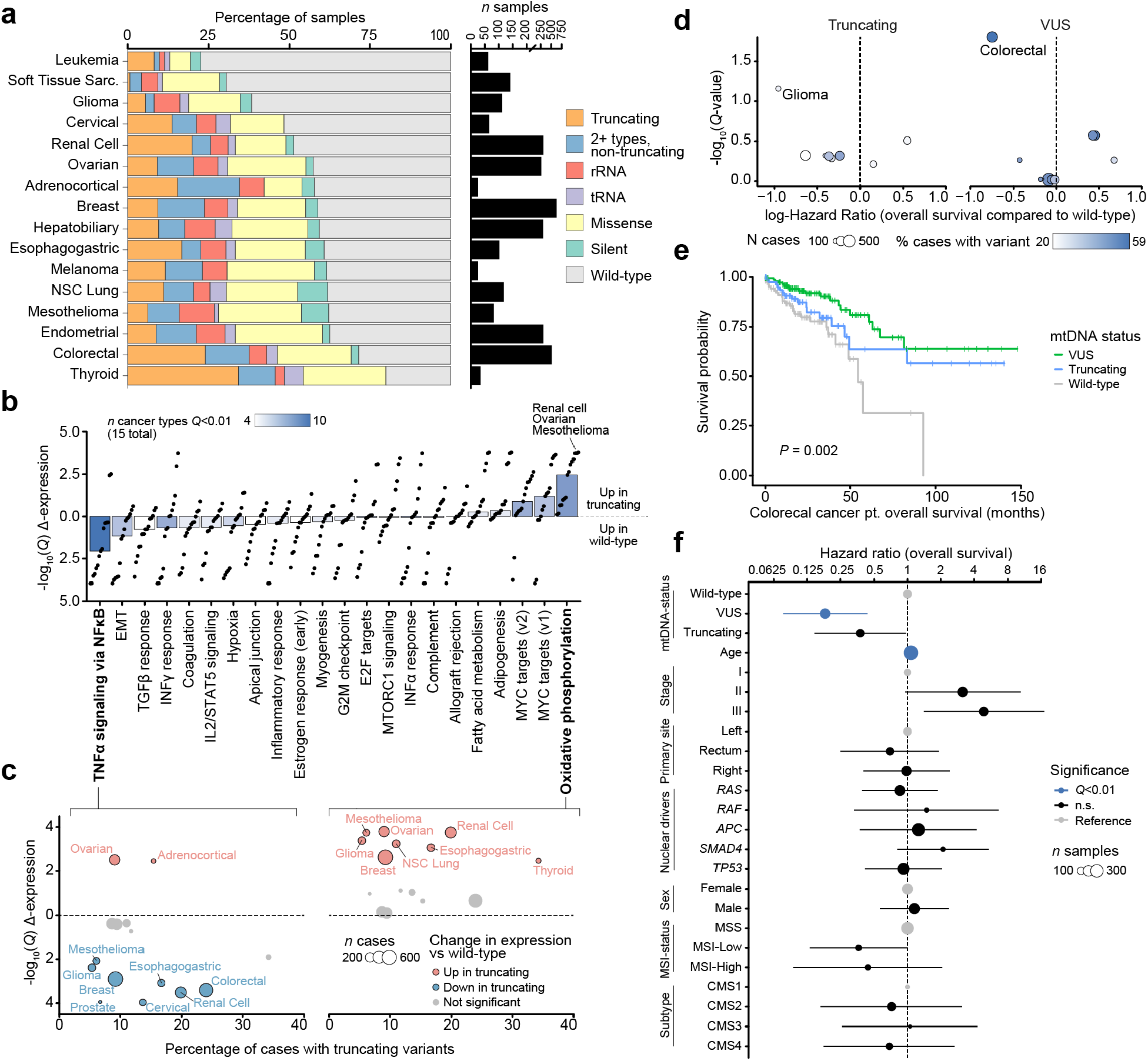
Mitochondrial genotypes associate with transcriptional and clinical phenotypes. **a)** Percentage of well-covered tumors with different types of somatic mtDNA variants per cancer type. Right, number of well-covered samples per cancer type. **b)** Differential expression of mSigDB Hallmarks genesets, between samples with truncating mtDNA variants and those with no nonsynonymous somatic mutations (*i.e.* “wild-type” samples). Differential expression is quantified by directional −log_10_(*Q*-value): greater than 0 denotes up-regulation in samples with truncating variants, below 0 denotes up-regulation in wild-type samples. Each dot is a single cancer type’s level of dysregulation of that geneset. Bars show the median level of dysregulation across 15 cancer types; bar shading shows the number of cancer types with significant dysregulation (*Q*<0.01) in either direction. **c)** Differential expression of TNFα via NFκB Signaling (left) and Oxidative Phosphorylation (right) genesets in individual cancer types. X-axis shows the overall proportion of samples of each cancer type with truncating variants; Y-axis matches the Y-axis in b). Dot width denotes number of well-covered samples for each cancer type. **d)** Effect size and statistical significance of mtDNA truncating variants (left) and VUSs (right) on overall survival among individual cancer types. Effect sizes (quantified as log-hazard ratios) are from univariate Cox proportional-hazards models run for each cancer type independently. *Q*-values are adjusted *P*-values from the model coefficients for each cancer type. **e)** Kaplan–Meier plot showing difference in overall survival time among *n*=344 TCGA colorectal cancer patients with somatic VUSs (*n*=152), truncating variants (*n*=84), or no nonsynonymous mutations (*i.e.* wild-type, *n*=108). **f)** Multivariate analysis of the effect of mtDNA variants on overall survival time among *n*=344 TCGA colorectal cancer patients (stage 1-3). Truncating variants and VUSs are each compared to wild-type samples, while controlling for known prognostic clinical and genomic covariates using a Cox proportional-hazards model. Hazard ratios for each covariate are shown on a log-scale, error-bars are 95% confidence intervals from the Cox proportional hazards regression. Point size indicates the number of samples with the associated covariate value (except for Age, which was coded as a continuous variable, and therefore the size corresponds to the total number of samples). Blue points are statistically significant (*Q*-value < 0.01); black points not significant; gray points are reference categories and were not tested.

Truncating mtDNA mutations approaching homoplasmy (>90% heteroplasmy) were identified in nearly all cancer types, despite the tendency for stromal and immune cell infiltration to suppress apparent tumor cell heteroplasmy, suggesting that even cancers in which mtDNA mutations are uncommon may still contain rare instances of individual tumors with highly mutant mitochondria.

In renal and thyroid tumors, truncating mtDNA mutations have historically been associated with the development of oncocytic neoplasia, whereby tumor cells accumulate dysfunctional mitochondria ^30,31^. That truncating mutations induce a morphologically similar response in two different tissue lineages suggests that cells may adopt a lineage-agnostic adaptation to the presence of a truncating mutation. To evaluate if truncating mutations induced functionally similar consequences across different tumor lineages, we compared the gene expression profiles of tumor samples with truncating mtDNA variants to tumor samples with wild-type mtDNA (harboring no nonsynonymous somatic mutations in protein-coding or RNA genes, see **Methods, Fig. 2f**). In half of all cancer types, tumors harboring truncating mutations exhibited a conserved expression program characterized by upregulation of genes associated with oxidative phosphorylation and downregulation of genes associated with TNFα via NFκB signaling (**Fig. 4b** and **Supplementary Fig. 4b**). Critically, these expression programs were evident in cancer types such as glioma and mesothelioma, where the proportion of samples with a truncating variant was comparatively low (**Fig. 4c**). These data suggest that, even in cancer types where mtDNA mutations are rare, truncating mtDNA mutations produce similar phenotypic outcomes.

Given that the hotspot *MT-ND1^R25Q^* exhibited an expression program resembling truncating variants, we investigated the generic transcriptional consequences of mtDNA VUSs (see **Methods**). Compared to truncating variants, fewer genesets demonstrated lineage-agnostic changes in samples with VUSs. As with truncating variants, the most upregulated geneset in VUS-harboring tumors was Oxidative Phosphorylation (increased in 5/18 cancer types) (**Supplementary Fig. 4c**), but the magnitude of this enrichment was attenuated relative to truncating variants. Notably, several cancer types, such as colorectal cancer, demonstrated a lineage-specific pattern of gene expression changes, suggesting that mtDNA VUSs are capable of eliciting a phenotype in specific cancer types.

To examine the translational value of mtDNA genotype, we determined the association between mtDNA mutation status and clinical outcome (overall survival) across cancer types. Using univariate Cox proportional-hazards regression, for each cancer type we determined the effect size and significance of both mtDNA truncating variants and VUSs compared to samples with no somatic mtDNA variants (wild-type). Colorectal cancer demonstrated the largest (by effect-size) significant association between overall survival time and mtDNA genotype (colorectal patients with VUSs had a hazard ratio of 0.47 (95%CI 0.03-0.75) compared to those with wild-type mtDNA, *Q*-value=0.02, Cox proportional-hazards regression) (**Fig. 4d**). Notably, VUSs in colorectal cancer also associated with a unique transcriptional down-regulation of multiple genesets including TNFα via NFκB, Hypoxia and Complement (**Supplementary Fig. 4c**, **Fig. 3j**), further suggesting a cryptic phenotype of these variants in affected tumors. We additionally observed a weak association between mitochondrial genotype and underlying molecular subtype ^32^, with some enrichment of mtDNA mutations in the canonical subtype CMS2 of colorectal tumors(**Supplementary Figure 4d**). We therefore further evaluated if mtDNA mutations may be prognostically meaningful in colorectal cancer, using a multivariate analysis to control for known prognostic clinical and genomic covariates. Among 344 stage 1-3 colorectal cancer patients, the presence of mtDNA alterations was significantly associated with better overall survival compared to wild-type samples (*P*=0.002, Kaplan–Meier test), with patients whose tumors harbored VUSs having the best prognosis, and those with truncating variants having an intermediate improvement (**Fig. 4e**). This association remained significant after controlling for clinically-relevant prognostic covariates (*i.e.* age, cancer stage, primary site, MSI-status, consensus molecular subtype and the presence of established nuclear-encoded genomic driver mutations) ^32,33^ in a multivariate analysis. VUSs again had a significantly protective association compared to wild-type (Hazard ratio=0.18, 95%; CI: 0.08-0.44; *Q*-value=0.001, Cox proportional-hazards model); truncating variants had an intermediate effect (HR=0.38, 95% CI: 0.15-0.97; *Q*=0.18) (**Fig. 4f**). These data together suggest that somatic mtDNA mutations are associated with a clinically and molecularly-distinct class of colorectal tumors, and that the functional consequence of an mtDNA mutation is a determinant of its clinical significance.

## Discussion

Although recent evolutionary data suggests that mtDNA mutations may be under positive selection in cancers of the kidney and thyroid ^5^, the broader significance of somatic mtDNA mutations in cancer remains a point of confusion and debate. Drawing inspiration from analyses describing hotspots of somatic mutations in the nuclear DNA of tumors, we studied the recurrence of mutant mtDNA alleles. The discovery that OXPHOS complex shapes mtDNA mutation patterns in a manner that produces mutation hotspots, in connection with orthogonal data on the structural consequences, transcriptomic effects and clinical significance of these alleles in patients with germline mtDNA disease, supports the hypothesis that mitochondrial respiration is perturbed across many tumors.

Our results indicate that OXPHOS complex, tissue lineage, and mutation consequence collectively shape the incidence and putative function of mtDNA mutations. Whereas previous studies have demonstrated localized regions of mtDNA with elevated somatic mutation rate in tumors, these works have generally been underpowered to probe phenotypic differences between alleles. Our data reveal that truncating mutations preferentially impact complex I, and that non-synonymous mutations of all classes are depleted in complex V. This suggests that cancer cells can better tolerate, or perhaps even utilize, loss of complex I and the associated metabolic consequences (e.g. NAD+:NADH changes), whereas loss of capacity for ATP synthesis through complex V mutations appears to be negatively selected against. That CIII demonstrates elevated rates (relative to other complexes) of missense mutations, but not truncating mutations, is consistent with its essential role in ubiquinol oxidation and suggests that weak disruption of CIII is preferential for clonal expansion in tumor cells ^21^. Whether truncating mutations in CIII and CIV promote different phenotypes in cancer cells relative to complex I loss warrants further investigation.

There is substantial evidence that in particular subtypes of thyroid and kidney cancer, mtDNA mutations are the root cause of metabolic adaptations and morphological (oncocytic) changes associated with suppression of mitochondrial respiration ^34^. What remains unclear is how to extrapolate the function of truncating mutations in otherwise essential mtDNA genes to cancer types where oncocytic tumors are rarely if ever observed but the fraction of samples harboring these mutations is nevertheless substantial (*e.g.* colorectal cancers). Critically, our transcriptional data suggests that, even in cancer types where truncating mtDNA mutations are rare, they nevertheless promote a transcriptional program characterized by increased expression of OXPHOS genes and downregulation of innate immune pathways. Because homoplasmic loss of any gene in the mtDNA necessarily cripples the cell’s ability to respire and disrupts dependent metabolic pathways, these findings suggest that pathogenic and high heteroplasmy mtDNA mutations potentially render a large fraction of tumors vulnerable to a metabolic therapeutic intervention.

## Methods

### Tumor and normal sample sequencing cohorts

Tumor and matched normal sequencing data for TCGA samples were obtained from the GDC Data Portal (https://portal.gdc.cancer.gov/). Briefly, all tumor and matched-normal barcodes included in the MC3 MAF ^35^ (https://gdc.cancer.gov/about-data/publications/pancanatlas) file were converted to UUIDs using the TCGAutils R package (v1.9.3), and these UUIDs were queried for whole-exome sequencing BAM files sliced for chrM using the GDC API. We then queried the GDC Data Portal for RNA-Sequencing BAM files for TCGA tumors already with whole-exome sequencing data. This process yielded paired tumor and matched-normal whole-exome sequencing BAMs for 10,132 TCGA patients, of which 9,455 had additional RNA-sequencing data. In addition to the raw sequencing data for TCGA samples from which we called mtDNA mutations (see: Calling mitochondrial variants), we additionally obtained somatic mitochondrial mutation calls for 2,836 whole-exome sequenced tumors from ICGC/PCAWG ^3^, of which 885 also had TCGA sequencing data. Nuclear somatic mutations for TCGA samples were obtained from the MC3 MAF, subset for the samples for which mtDNA whole-exome sequencing BAMs were available. Finally, mtDNA mutation calls for 195,983 normal samples were obtained from the HelixMTdb cohort of sequenced saliva samples from healthy individuals ^14^.

### Annotating mtDNA regions included in our analysis

Each mitochondrially-encoded gene’s name, start/end positions and DNA strand was obtained from Biomart for human reference genome GRCh38 (release 95). Subsequently, each mtDNA position (1-16569) was annotated with its associated genetic information. Any mtDNA positions located at the overlap of two genes were annotated only as associated with whichever gene started first in numerical genomic position. Variants in non-genic mtDNA regions were excluded in our analyses. To this end, we excluded any variants in the mtDNA Control Region (positions 1-576, 16,024-16,569) as well as 89 other non-genic positions. We similarly excluded variants in hypermutated regions of mtDNA, including 302-316, 514-524, and 3,106-3,109). Following these measures, the genomic length of mtDNA retained in our analyses was 15,354bp. (The complete list of 16,569 mtDNA positions and their annotated reasons for exclusion is provided in **SI table 1**.)

### Calling mitochondrial variants

Mutations to the mitochondrial genome were obtained from variants called by both of two independent variant-calling pipelines. In the first pipeline, Mutect2 (GATK v4.1.2.0) ^36^ was used to call variants in chrM in tumor and normal samples individually, the results of which were subsequently intersected to obtain variants called supported in a given patient’s tumor and matched normal samples. Briefly, Mutect2 was run in mitochondrial-mode for each patient’s tumor and normal sample independently against human reference genome GRCh38 (with minimum base quality-score 20, minimum mapping quality 10, aggressive pcr-indel model, and other standard quality control arguments for paired-end reads). Artifacts were subsequently removed using GATK *FilterMutectCalls* (GATK v4.1.2.0) ^36^, and multi-allelic sites were split into individual variants using the *norm* function from bcftools (v1.9) ^37^. The resulting tumor and normal VCF files were then merged using gatk *HaplotypeCaller* (GATK v4.1.2.0) ^36^, to annotate variants in the tumor VCF with their coverage in the normal sample. The resulting VCF was converted to a MAF file using vcf2maf (v1.6.17, https://github.com/mskcc/vcf2maf). Finally, variants from the generated MAF file were then filtered out unless the variant allele was supported at least one read in both forward and reverse directions. In the second pipeline, samtools mpileup (v1.9) ^11^ was used to generate a pileup file using variant-supporting reads with minimum mapping quality 20 and base alignment quality 10. Reads failing quality checks or marked as PCR duplicates were removed. Variants were required to contain at least 2 variant-supporting reads in the forward and reverse direction. In each pipeline, variants were additionally filtered to ensure ≥ 5% variant allele frequency in the tumor, and tumor coverage ≥ 5 reads. Variants identified by both pipelines were retained for further analysis. In rare cases, multiple indels were called in a sample within a homopolymeric region (single-nucleotide repeats of 5 or more basepairs), with distinct alt-read counts and VAF values, and identical read-depth values. These multiple indels were collapsed to a single representative indel call. Briefly, using the Mutect2 variant calls, whichever indel had the highest VAF in the tumor sample was taken as the representative indel. The count of alt-reads in both tumor and normal were replaced with their corresponding summed counts across the original multiple indels, and the VAFs in both tumor and normal were re-calculated from the new summed alt-read counts divided by the original read-depth. Finally, mutations were classified as of somatic origin according to the following criteria: Non-truncating variants (that is, all variant classifications other than nonsense mutations and frame-shift indels) were classified as somatic if the matched normal sample had a minimum coverage of 5 reads and 0 normal reads called the alternate allele. Truncating variants in the tumor sample were assumed to be of somatic origin. All other variants were not classified as somatic and excluded from this study.

### Nuclear mutational data and annotation

Somatic mutations in nuclear-encoded cancer-associated genes for TCGA samples were obtained from the PanCanAtlas MC3 MAF file. Mutations in this file were subset for those among the 468 genes on the MSK-IMPACT clinical sequencing panel ^13^. The MAF file was annotated for known, likely, and predicted oncogenic driver mutations using the MAF-Annotator tool provided by OncoKB ^38^ (https://github.com/oncokb/oncokb-annotator). Mutations annotated by OncoKB as “Oncogenic”, “Likely Oncogenic” or “Predicted Oncogenic”, previously determined cancer hotspot mutations ^15,16^, or truncating variants to tumor suppressor genes (*i.e.* frame-shift indels, splice-site and nonsense mutations) were classified as potential driver alterations.

### Calculating tumor mutational burden in mtDNA or nuclear DNA

Tumor mutational burden (TMB) was calculated for cohorts of tumors subset for various genomic regions, including: 1) individual mitochondrial- or nuclear-encoded genes; 2) mtDNA genes grouped by OXPHOS complex I, III, IV, or V; 3) the entire mitochondrial genome (excluding non-genic and polymorphic regions); 4) a set of known nuclear-encoded tumor suppressor genes; and 5) a set of known nuclear-encoded oncogenes. In each case, TMB was calculated as the total number of somatic mutations among the relevant collection of tumors divided by the total genomic length sequenced in these tumors (in Mbps). For TMBs calculated from mutations called in off-target sequencing data (*i.e.* mtDNA variants in TCGA samples), the total genomic length sequenced was the number of the genomic positions with sufficient coverage to call somatic variants (5+ read coverage in both tumor and normal sample), summed across all samples. For TMBs calculated from targeted regions (nuclear DNA; mtDNA in PCAWG samples), the total genomic length sequenced was the length of the targeted region (entire gene for mtDNA, exonic regions for nuclear DNA) multiplied by the number of samples. Error bars for TMBs were calculated as 95% Poisson exact confidence intervals for rates, using the total number of mutations as the count of events, and the genomic length sequenced in Mb as the time at risk.

### Identifying hotspot positions for mitochondrial variants

We identified mtDNA positions with statistically recurrent single-nucleotide variants (SNVs) by comparing the observed proportion of mutations at an individual position (out of the total number of mutations acquired in its gene) to a rate of mutations at the position expected by chance with a one-sided binomial test. The probability for SNVs at each position of a gene *P_pos,gene_* was modeled as a bernoulli trial, where the likelihood of a mutation arising at a given position by its mutability relative to the mutability of all other bases in the gene: 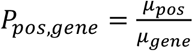. Consistent with previous work ^15^, we estimated the mutability for each position as a function of its trinucleotide context. That is, for each position, it’s mutability *μ*_*pos*_ was calculated as the count of SNVs matching the trinucleotide context of the position of interest *S*_*pos*_, out of the total count of SNVs anywhere in the mitochondrial genome *S*_*total*_ (after excluding the control region and other blacklisted regions). Due to the highly strand-specific mutation signatures we observed for SNVs in mtDNA (**Supplementary Fig. 1c**), we used the complete set of 64 unique trinucleotides in order to retain this information when calculating the mutability for each position, rather than collapsing the central nucleotide to C or T resulting in the conventional 32 unique trinucleotides. As the proportion of patients for which a given position had sequencing coverage in paired tumor and normal samples linearly affects the likelihood of observing a somatic mutation at the position, the mutability of a position was adjusted to control for this by multiplying it by the ratio of the number of samples with paired tumor-normal sequencing coverage at the position *C*_*pos*_ out of the total number of samples *N_samples_*, so that 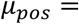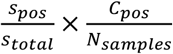. The mutability associated with the gene was calculated as the sum of each position’s trinucleotide mutability. Therefore, for a gene *L* basepairs in length: 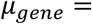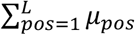. The final parameter for the binomial test (i.e. the likelihood for a mutation in a gene to arise at the given position by chance) was therefore 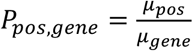. Each position mutated in 5 or more samples in each gene was subsequently tested for statistically enriched mutations by comparing its observed number of mutations out of the total number of mutations in the gene to this binomial parameter using a right-tailed binomial test. The full list of generated *P*-values across all genes were then corrected for multiple hypothesis testing.

### Homopolymer hotspots for indels

To identify homopolymer regions with statistically enriched rates of insertions and deletions (indels), we modeled the proportion of samples with indels across all homopolymers as a function of the homopolymer region’s width (i.e. the number of repeated nucleotides, from 5-8). To this end, all single-nucleotide repeats of 5 or more basepairs were identified in the mitochondrial reference genome, resulting in N=73 unique homopolymer loci in whitelisted coding mtDNA. We then modeled the fraction of frame-shift indels across 73 homopolymers observed to arise at a specific homopolymer locus *h* as a binomial process dictated by the length of the homopolymer *l_h_* divided by the summed length of all homopolymers, such that the expected likelihood of a frame-shift indel arising at a homopolymer by chance is given by: 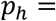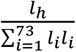. We then tested each homopolymer locus for enriched mutations with a one-sided binomial test. That is, for each homopolymer locus, the number of bernoulli trials was the number of samples with complete sequencing coverage for the homopolymer region and two flanking basepairs; the number of successes was the number of samples with frame-shift indels at (or immediately adjacent to) the given homopolymer, and the fraction of successful trials was compared to the expected probability *p_h_*.

### Hotspot positions in tRNA cloverleaf structure

Positions of the tRNA cloverleaf secondary structure were individually tested for an enriched rate of SNVs at the equivalent aligned positions of the 22 mitochondrially-encoded tRNAs. A map of genomic positions in mitochondrial tRNAs to cloverleaf structure positions was provided by Mitotip ^39^ (https://github.com/sonneysa/MitoTIP/blob/master/Output/tRNA%20data%20and%20scoring_scored.xlsx) and used to assign SNVs at tRNAs to structural positions. Under the null hypothesis that mutations accumulate at structurally-aligned positions randomly, the proportion of SNVs aligning to a specific position in the tRNA cloverleaf should be approximately equal to the number of times the aligned position was sequenced at a sufficient depth in both tumor and matched normal samples to call somatic mutations, out of the total number of tRNA basepairs sequenced at sufficient depth across all samples at all structural positions. Therefore for a given position of the tRNA cloverleaf structure *p*, the number of SNVs observed across all tRNAs at this aligned position *t_p_* out of *T* SNVs across all positions of all tRNAs was tested for enrichment using a one-sided binomial test, compared to an expected rate equal to the number of tRNA bases aligned to this position sequenced at sufficient depth *b_p_* out of *B* tRNA bases sequenced at sufficient depth across all positions of all tRNAs.

### Classifying sample mtDNA variant status

Each tumor sample was classified according to the presence and type of its somatic mitochondrial variants. Because gaps in sequencing coverage may make existing variants undetectable and result in the incorrect classification of such samples as “wild-type” for somatic variants, we only attempted to classify samples with sequencing coverage in both tumor and matched normal of at least 90% of the included region of mtDNA (referred to as “well-covered” throughout). Furthermore, given the high incidence of truncating indels we observed at 6 hotspot loci, we additionally required that these 6 loci were sequenced at sufficient coverage in the tumor sample, to ensure that samples potentially harboring recurrent indels would be excluded and not misclassified. Samples not meeting either of these conditions were classified as having ‘Unknown’ mtDNA mutation status. The remaining samples were then classified according to a decision tree as follows: Samples with any protein-truncating variants were classified as ‘Truncating’; remaining samples still unclassified with multiple mtDNA variants of different types (among missense, rRNA, and tRNA variants) were classified as ‘2+ non-truncating types’; remaining samples with tRNA mutations were classified as ‘tRNA’; remaining samples with rRNA mutations were classified as ‘rRNA’; remaining samples with non-truncating, non-synonymous protein-coding mutations as ‘missense’; remaining samples with silent mutations as ‘Silent’; and finally samples still unclassified were classified as ‘wild-type’. This logic prioritizes minimizing annotation bias over conserving sample size, in order to meaningfully compare the incidence of different variant types across samples. However, in our analysis of the effect of mtDNA variants on differential gene expression or survival, we modified the logic to prioritize conservation of sample size. To this end, in RNA-Seq and survival analyses, samples with any observed truncating variants were classified as truncating, regardless of their sequencing coverage.

### Testing genesets for transcriptional dysregulation due to mtDNA variants

A matrix of estimated gene expression counts (RSEM values normalized to correct for batch effects) for TCGA samples was downloaded from the TCGA PanCanAtlas ^35^ supplemental data (http://api.gdc.cancer.gov/data/3586c0da-64d0-4b74-a449-5ff4d9136611). Gene expression estimates were rounded to integer values, and subsequently genes with zero estimated counts in all samples were removed, as were genes with unknown gene symbols. To evaluate differentially expressed genes between two groups of samples with different mtDNA variant type (i.e. truncating vs wild-type samples colorectal samples), the rounded gene expression matrix was subset for the relevant samples and input into the DESeq2 ^40^ package in R using the DESeqDataSetFromMatrix utility, along with a table of tumor sample barcodes with their associated mtDNA classification. Differentially expressed genes were tested and their log-fold change (LFC) values were shrunken using the apeglm ^41^ package. *P*-values for all genes tested were corrected for multiple-hypothesis testing with the Benjamini Hochberg method ^42^. The resulting data from this analysis were used to calculate a statistic for each gene equal to log_10_(*Q*-value)×sign(LFC). All genesets from the mSigDB Hallmark geneset collection ^43^ (v7.1) were then tested for significant up- or down-regulation based on this statistic for each gene using the fgsea package ^44^ in R, with a minimum geneset size of 10 genes, a maximum size of 500 genes, and 100,000 permutations.

### Annotating genomic and clinical covariates in colorectal cancer survival analysis

Clinical data for TCGA colorectal cancer patients including: overall survival time/status, AJCC pathologic tumor stage, age at diagnosis, sex, and tumor tissue site were obtained from the TCGA FIrehose legacy data on cbioportal (https://www.cbioportal.org/study/summary?id=coadread_tcga). Clinical data was subset for patients with sequencing data in the MC3 MAF. These data were then annotated with MSI status (MSS, MSI-low, MSI-high) based on published data for patients where this was available 45. AJCC Pathologic Tumor Staging data was collapsed into Stages I, II, III, IV, and Stage-IV patients were excluded. The tumor site was encoded as “Right-colon” if the primary site was: ascending colon, cecum, hepatic flexure, or transverse colon; or encoded as “Left-colon” for: descending colon, sigmoid colon, or splenic flexure. Patients with tumor tissue from the rectum were encoded as “Rectum” for their tumor site. The clinical data for each sample was then annotated for the presence of known or likely nuclear-encoded driver alterations in KRAS/HRAS/NRAS, BRAF, APC, SMAD4 and TP53 as based on mutation calls from the TCGA MC3 MAF ^46^ (see: Methods “Nuclear mutational data and annotation”). Each patient in the clinical data was then annotated as having a known/likely driver alteration in each of KRAS/HRAS/NRAS (grouped into RAS), BRAF, APC, SMAD4 or TP53. The complete multi-variate model use in the Cox proportional-hazards regression was therefore: Overall Survival ~ mtDNA-status + Age + Stage + Site + RAS + RAF + APC + SMAD4 + TP53 + Sex + MSI-status + CMS-type.

### Structural impact of *MT-ND1^R25Q^* variant on complex I

The structural impact of the *MT-ND1^R25Q^* variant was investigated using an electron-microscopy derived structure of mitochondrial CI in *mus musculus* (PDBID: 6G2J) ^29^. The UCSF Chimera software (v1.13.1) ^47^ was used to insert the R25Q mutation using the *swapaa* command. The ubiquinone binding tunnel was predicted using the CAVER Analyst (v2.0b) ^48^ software run on the wild-type PDB structure, starting from the side chain oxygen atom in *Ndufs2^Y108^*, and using a minimum probe radius of 1.4Å as described by the authors ^49^. Surface electrostatic charge for wild-type and mutant structures were determined using the APBS software ^50^ (http://server.poissonboltzmann.org/pdb2pqr) using default parameters, after subsetting the PDB structure for Mtnd1 (chain H), and converting the resulting PDB file to PQR using PDB2PQR ^51^. All structure visualizations were generated using UCSF Chimera.

### Statistical analyses and figures

All statistical analyses were performed using the R statistical programming environment (version 3.6.1). Protein structure figures were generated using UCSF Chimera, Kaplan-Meier plots and Cox proportional hazard forest plots were generated with the survminer library in R, ETC schematic (Fig. 1a) in Adobe Illustrator. All other figures were generated using the ggplot2 library in R. Unless otherwise noted, error bars for proportions are 95% binomial CIs calculated using the Pearson-Klopper method; error bars for rates (e.g. Mutations/Mb) are 95% Poisson CIs calculated with the pois.exact function from the epitools library in R. Unless otherwise noted, *P*-values for difference in proportions were calculated using Fisher’s exact tests or two-sample Z-tests, and for difference in rates using Poisson exact tests. *P*-values were corrected for multiple comparisons using the Benjamini-Hochberg method ^42^ and reported as *Q*-values when applicable.

## Data and code availability

All relevant data and R code are available on GitHub with instructions to execute the code and regenerate all figures (https://github.com/reznik-lab/mtdna-mutations).

## Acknowledgements

We thank the members of the Reznik and Taylor laboratories for discussion and support. We also thank Lydia Finley, Kivanc Birsoy, and Nicole Rusk for their feedback.

## Author contributions

ANG, PAG, and ER conceived the study.MK, WKC, KL, AAH, and BST assisted with genomic data analysis. ANG, PAG, and ER wrote the manuscript with input from all authors.

## Competing financial interests

The authors declare no competing financial interests

## Supplementary Materials

### Supplementary Tables

**Supplementary Table 1:** Table of mtDNA position 1-16,569 annotated with gene symbols, encoding strand, nucleotide, and exclusion criteria.

**Supplementary Table 2:** Table of mutation rates in mtDNA and nuclear cancer-associated genes.

**Supplementary Table 3:** Table of SNV hotspot positions and associated significance and annotations.

**Supplementary Table 4:** Table of homopolymer indel hotspots and associated significance and annotations.

**Supplementary Table 5:** Table of tRNA structural alignment hotspots and significance and annotations.

**Supplementary Table 6:** Table with mtDNA variants and mtDNA status classifications for all TCGA samples included in this study.

### Supplementary Figures

**Supplementary Fig. 1a:**
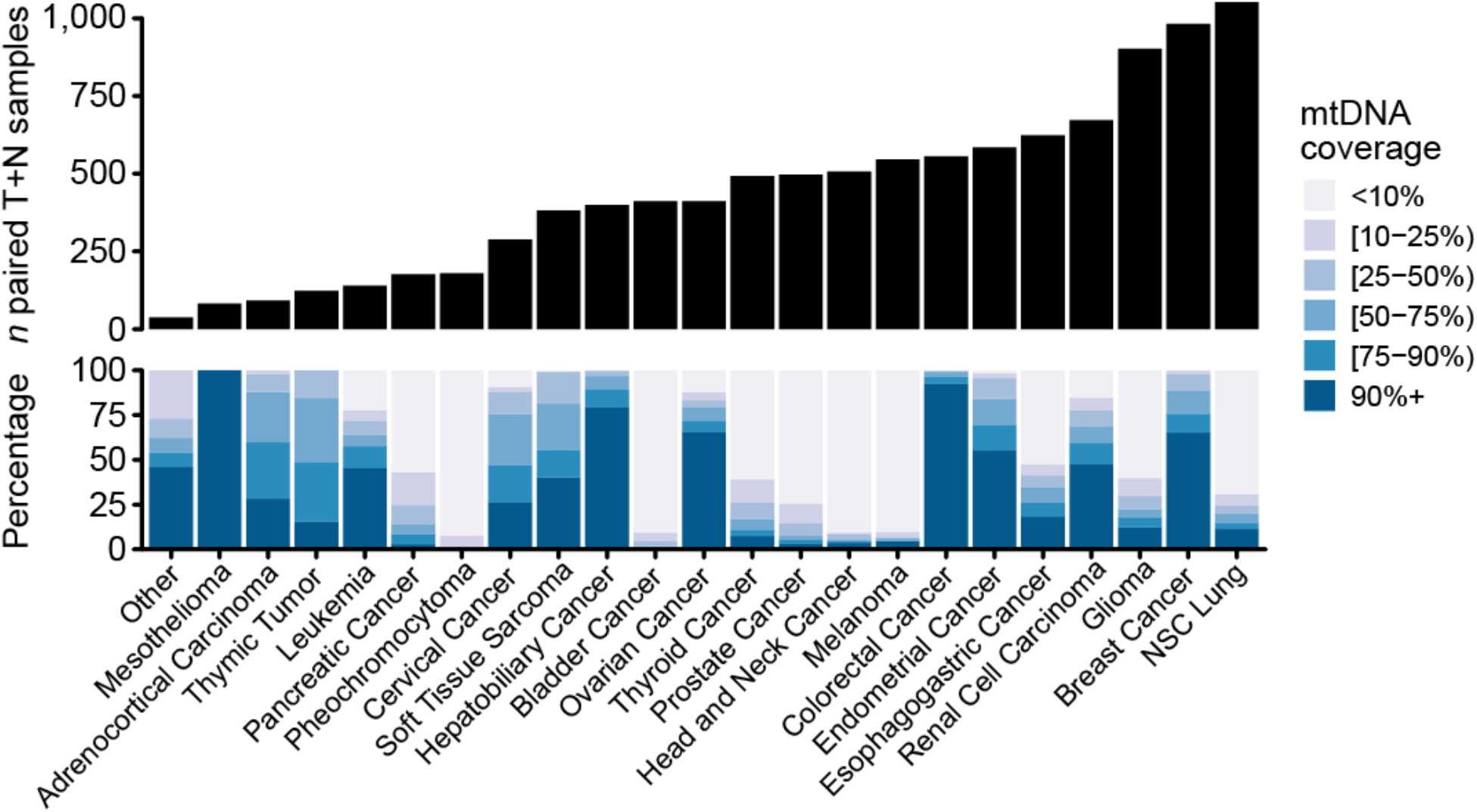
Distribution of cancer types in patient cohort. Top, number of tumor samples and paired matched normal samples per cancer type in this study. Bottom, the proportion of tumor and normal sample pairs each with ≥5 read coverage in the indicated percentage of genic regions of the mitochondrial genome (*e.g.* darkest blue indicates the percent of well-covered samples of the given cancer type.).

**Supplementary Fig. 1b:**
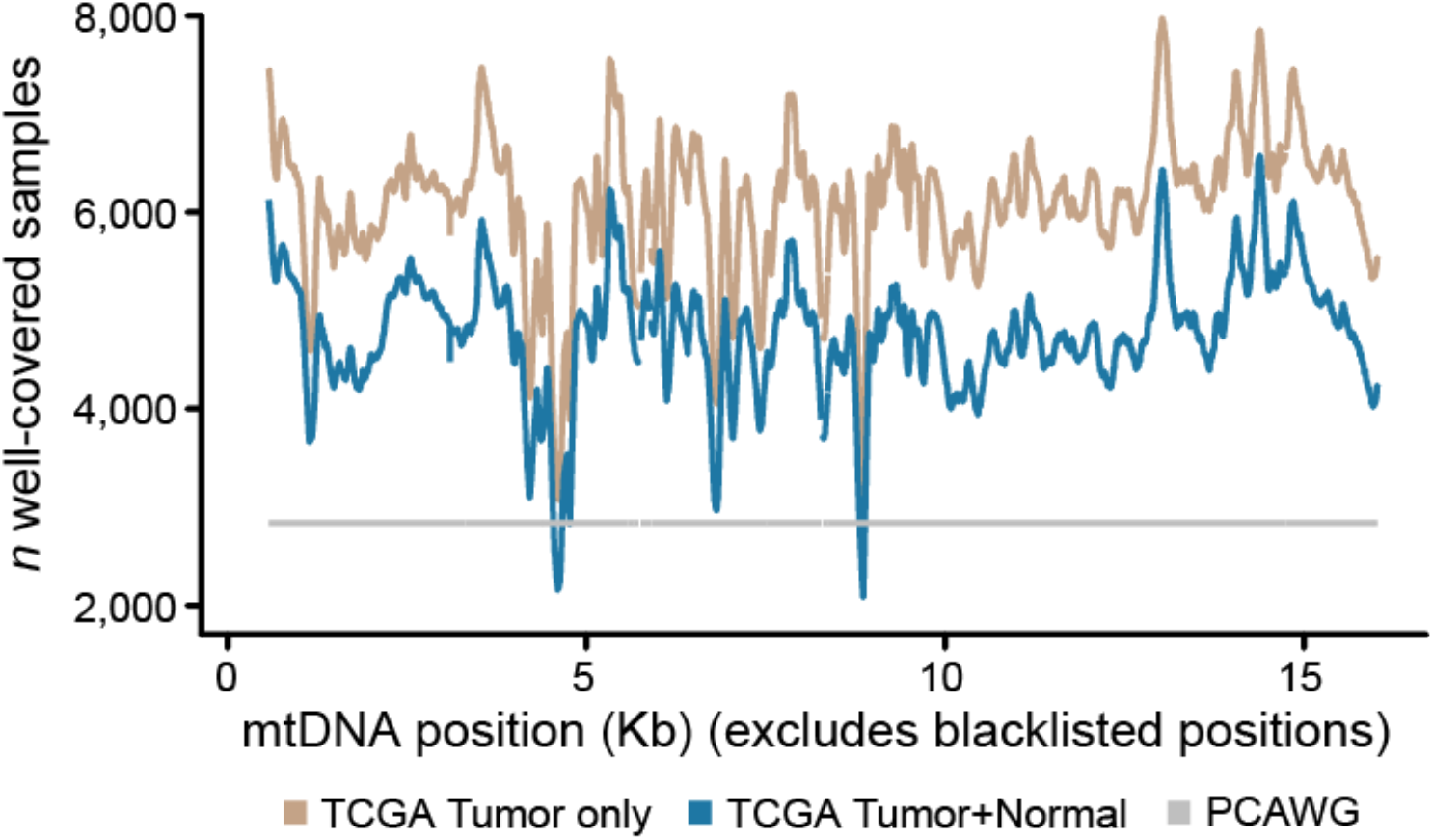
mtDNA coverage from off-target reads at each position. The number of samples for which the given mtDNA position was sequenced to adequate depth to call somatic variants. Brown, the number of samples using unpaired tumor-only data, applicable only for protein-truncating variants which were always assumed to be of somatic origin. Blue, the number using paired tumor and matched-normal data, applicable for all non-truncating variants which required evidence that the variant was absent in the matched normal to be confidently classified as of somatic origin. Gray, the number of whole-genome sequenced samples available from ICGC/PCAWG for comparison.

**Supplementary Fig. 1c:**
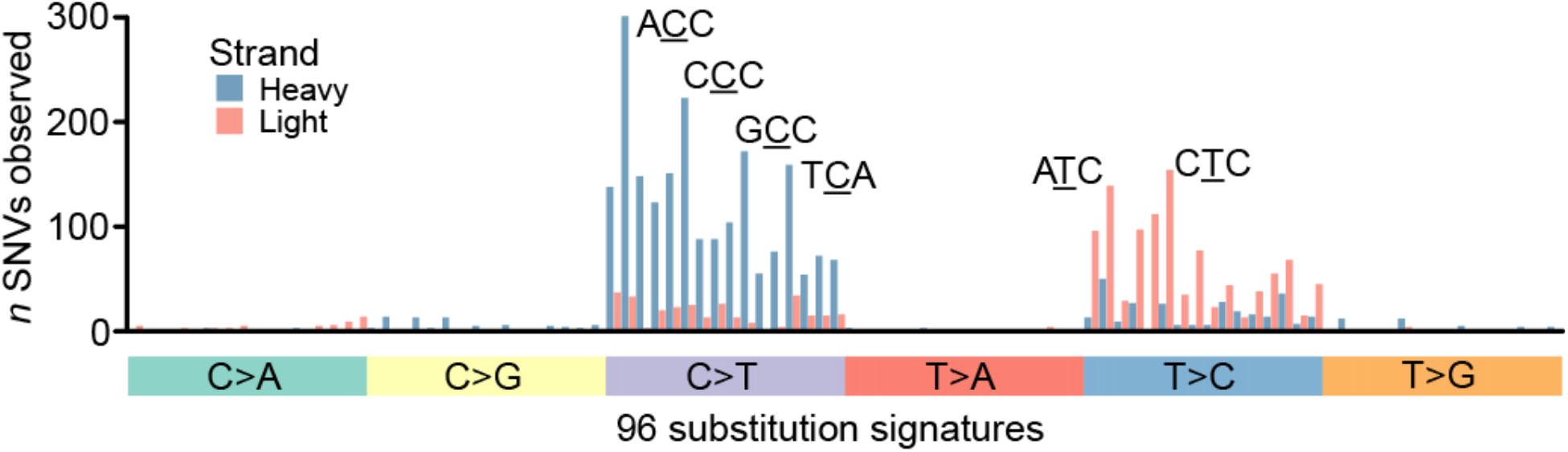
Strand-specific mutational signatures in our dataset. The frequency of somatic SNVs on the light or heavy mtDNA strand with each of the 96 possible mutational signatures with trinucleotide contexts (among *n* = 3,872 SNVs). Blue bars indicate the prevalence of mutational signatures for heavy-strand encoded SNVs (substitutions at C or T central nucleotides); red bars indicate those for light-strand encoded SNVs (substitutions at G or T nucleotides, which were standardized to their C or T complementary nucleotide). The most prevalent mutational signatures are labeled. The underlined central position is mutated with the single nucleotide substitution labeled in the tile below.

**Supplementary Fig. 1d:**
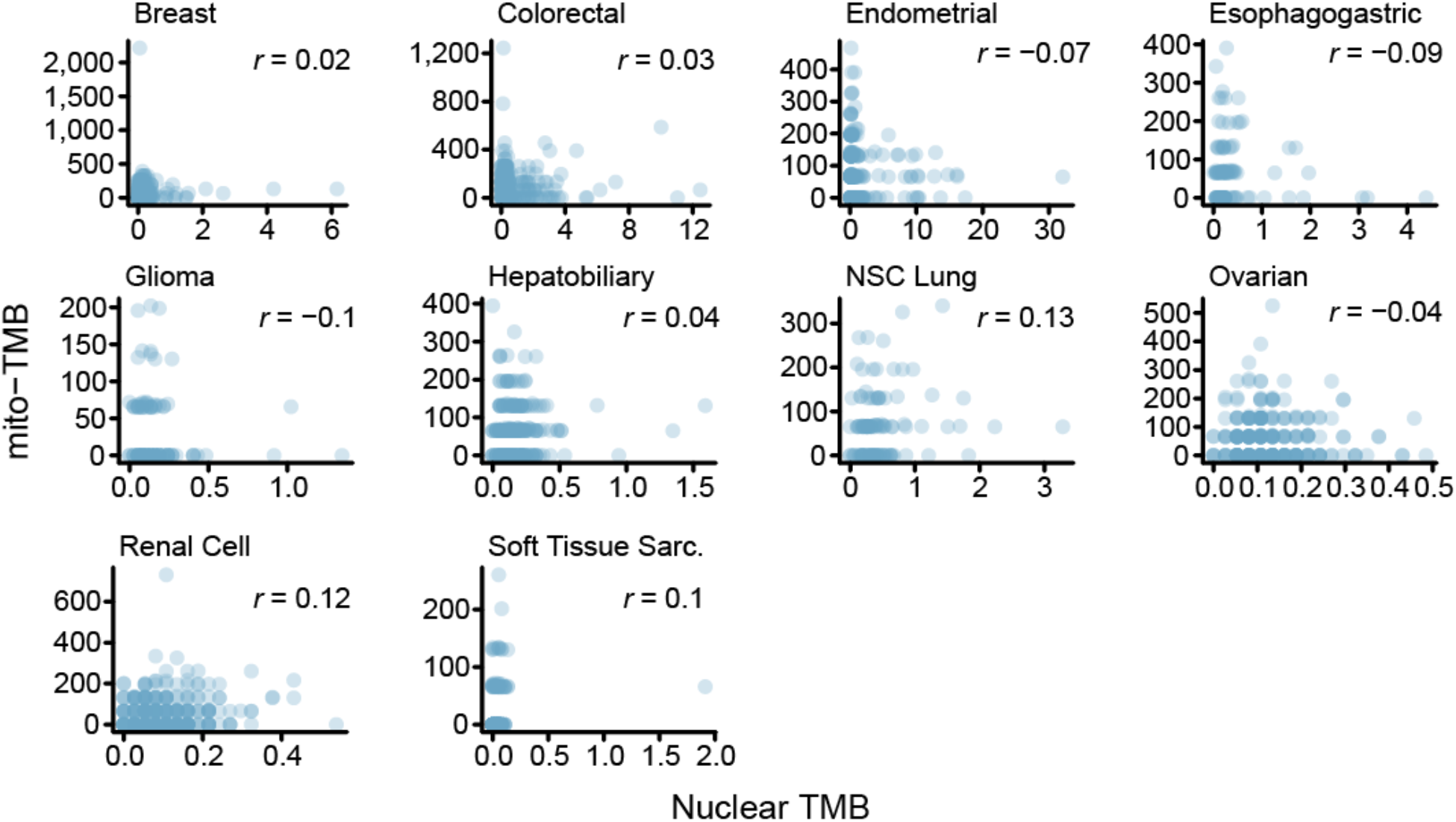
mtDNA mutation burden does not correlate with nuclear mutation burden within cancer types. Mitochondrial and nuclear tumor mutation burdens (TMB, mutations/Mb) are shown for each well-covered tumor, among cancer types with *n* ≥ 100 samples. Nuclear TMBs are calculated based on mutations to 468 cancer-associated genes and their total coding-sequence length. Pearson correlation coefficients *r* indicate no linear correlation between mitochondrial and nuclear TMBs were observed for any cancer type tested.

**Supplementary Fig. 1e:**
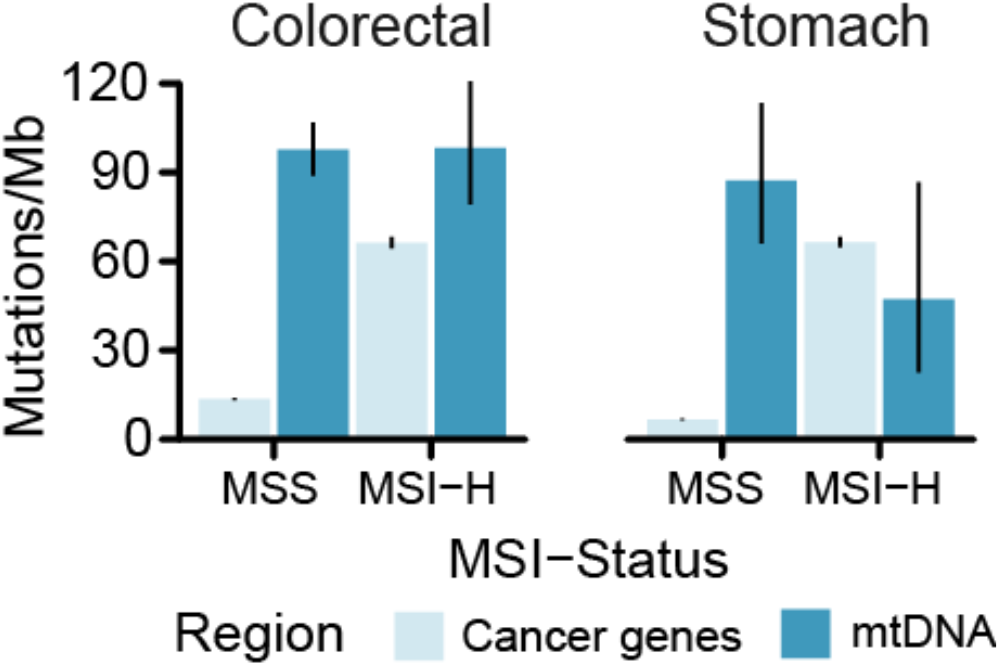
Microsatellite instability does not affect somatic mtDNA mutation rate. TMBs for somatic mtDNA mutations and mutations to cancer-associated genes are compared between microsatellite stable (MSS) and microsatellite unstable (MSI-High) tumors, for both (*n* colorectal cancer: MSI=65, MSS=318; *n* stomach adenocarcinomas: MSI=75, MSS=256). Although MSI-High tumors have elevated TMB for nuclear cancer genes, there is no effect on mtDNA TMB. Moreover, mtDNA TMB is similar to (or exceeds) that of nuclear cancer associated genes in both cancer types. Error bars are 95% Poisson exact confidence intervals.

**Supplementary Fig. 2a:**
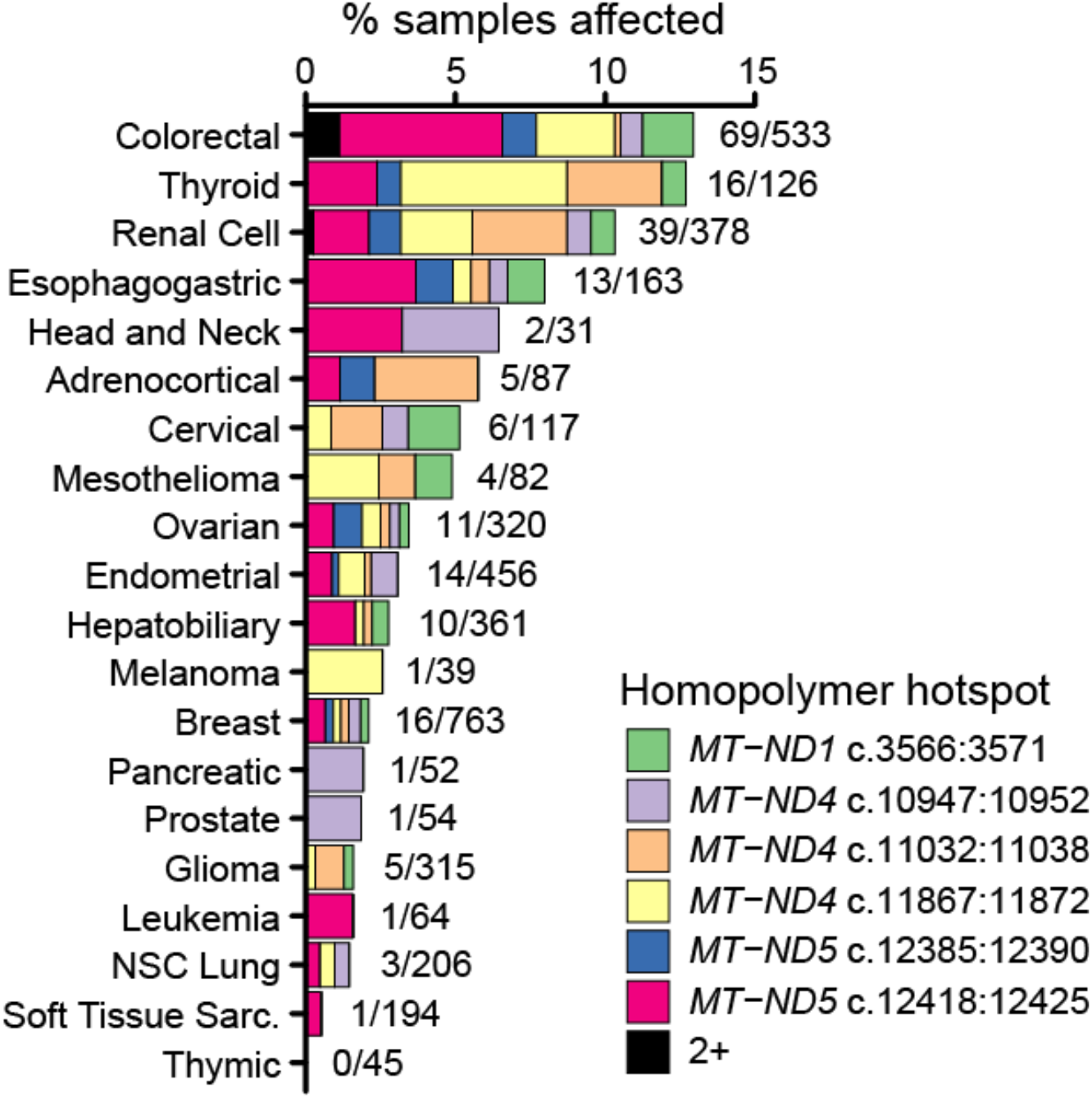
Prevalence of frame-shift indels at homopolymer hotspots across cancer types. Percentage of cases per cancer type with truncating frame-shift indels at any of 6 indel hotspot loci. Plotted cancer types had ≥ 20 well-covered samples (*n*=4,432 paired tumor and matched-normal samples total). Labels indicate the fraction of samples with any indels at homopolymer hotspot out of the total number of well-covered samples for the given cancer type.

**Supplementary Fig. 2b:**
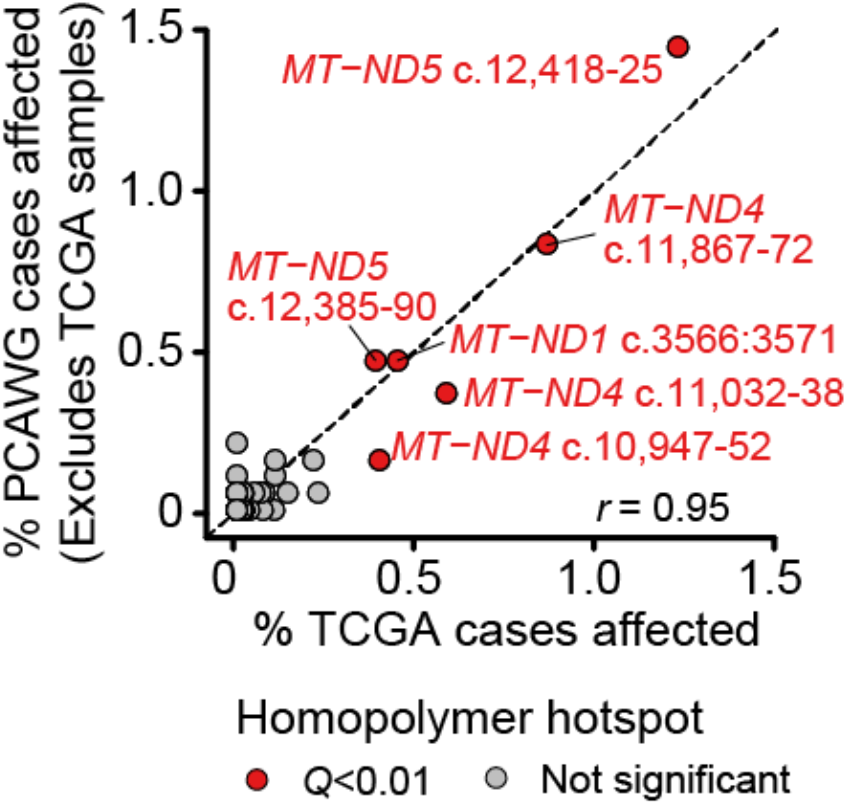
Validation of homopolymeric indel hotspot loci. The proportion of samples in TCGA (X-axis) or PCAWG (excluding samples also in TCGA, Y-axis) with frame-shift indels at 73 homopolymeric regions. The 6 indel hotspot loci are colored red and labeled. y=x is shown as a dashed line. Pearson correlation coefficient *r* as indicated.

**Supplementary Fig. 3c:**
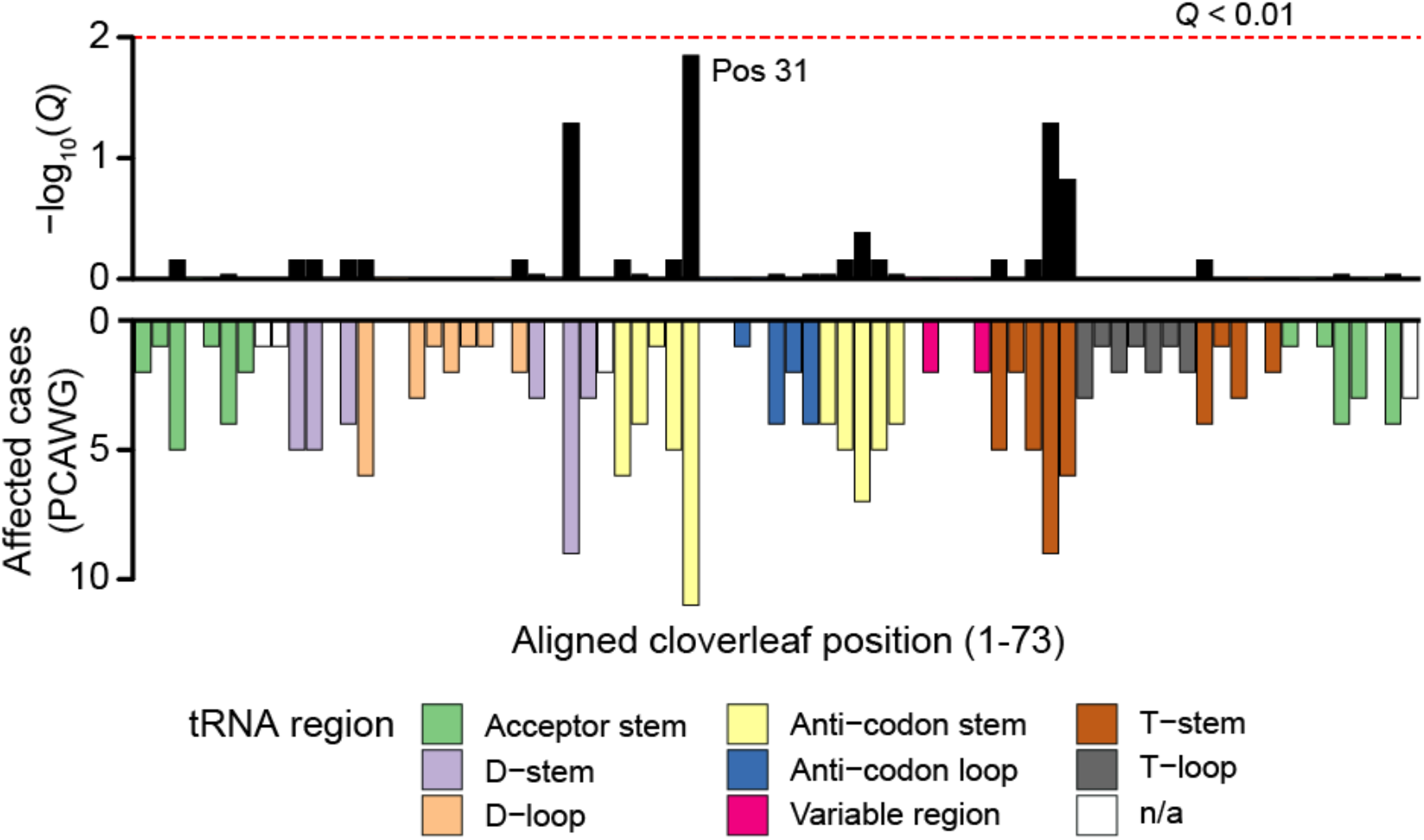
Validation of tRNA structural hotspots in PCAWG. The number of samples with SNVs in tRNAs at the indicated cloverleaf structural position, bottom; top, the statistical enriched of the given position for mutations. Position 31 *Q*-value=0.014, *n*=196 tRNA mutations among 1,951 PCAWG samples.

**Supplementary Fig. 4a:**
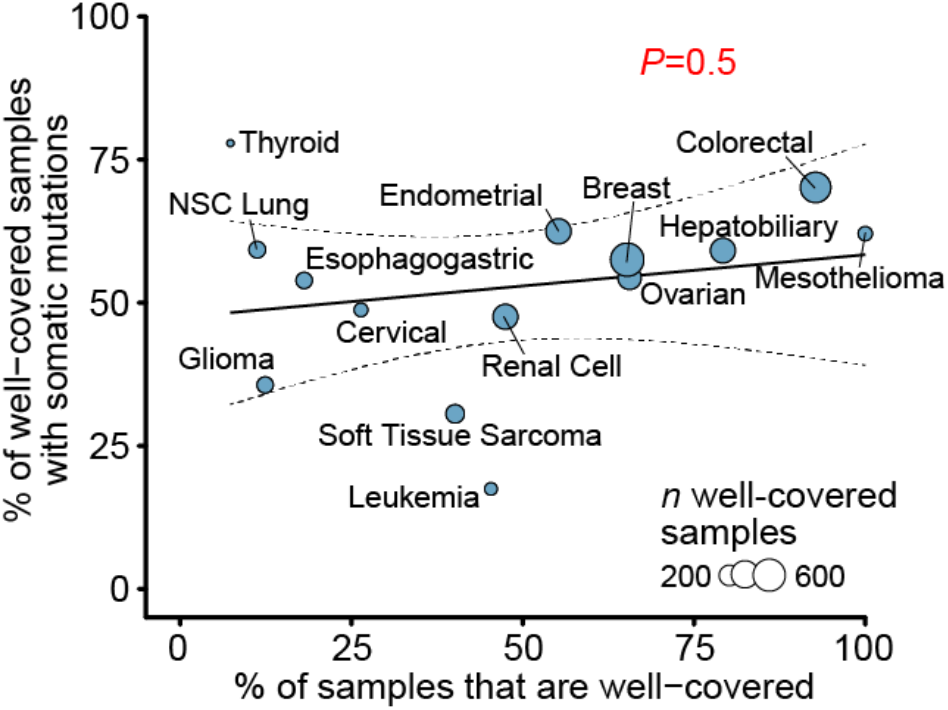
Proportion of samples with detectable mutations is not biased by cancer type sequencing coverage. There is no correlation between the fraction of well-covered samples in a cancer type and the proportion of well-covered samples with a detectable somatic mtDNA mutation. Cancer types with ≥30 well-covered samples shown, *P* value from linear regression.

**Supplementary Fig. 4b:**
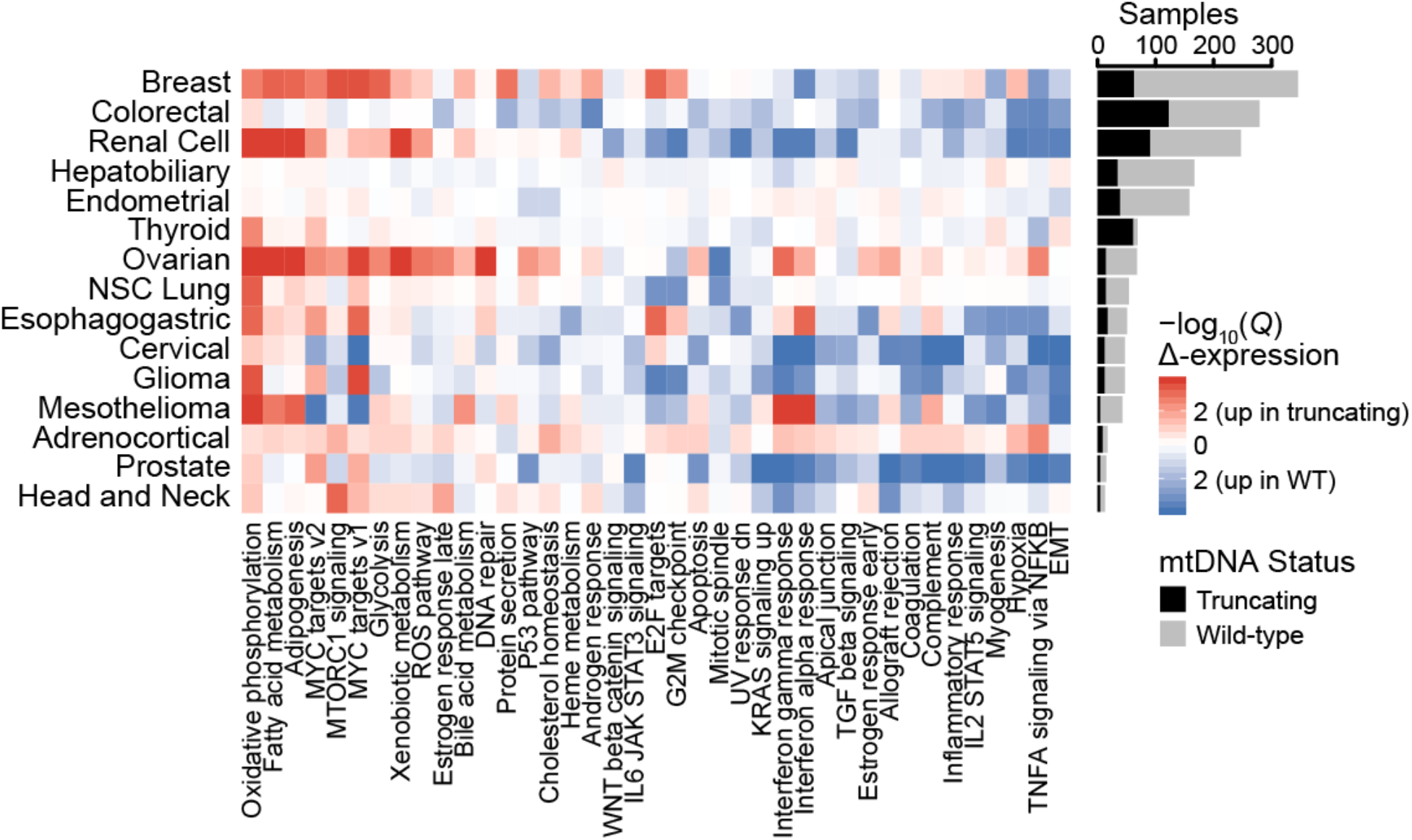
Transcriptional dysregulation attributed to truncating mtDNA variants. (Left) Heatmap shows directional significance of dysregulation of a given geneset in tumors with truncating variants among the given cancer type; −log_10_(*Q*-value) > 2 indicates significant up-regulation, < −2 indicates significant down-regulation. (Right) Histogram of wild-type samples and samples with truncating variants used to calculate differentially-expressed genes and dysregulated genesets.

**Supplementary Fig. 4c:**
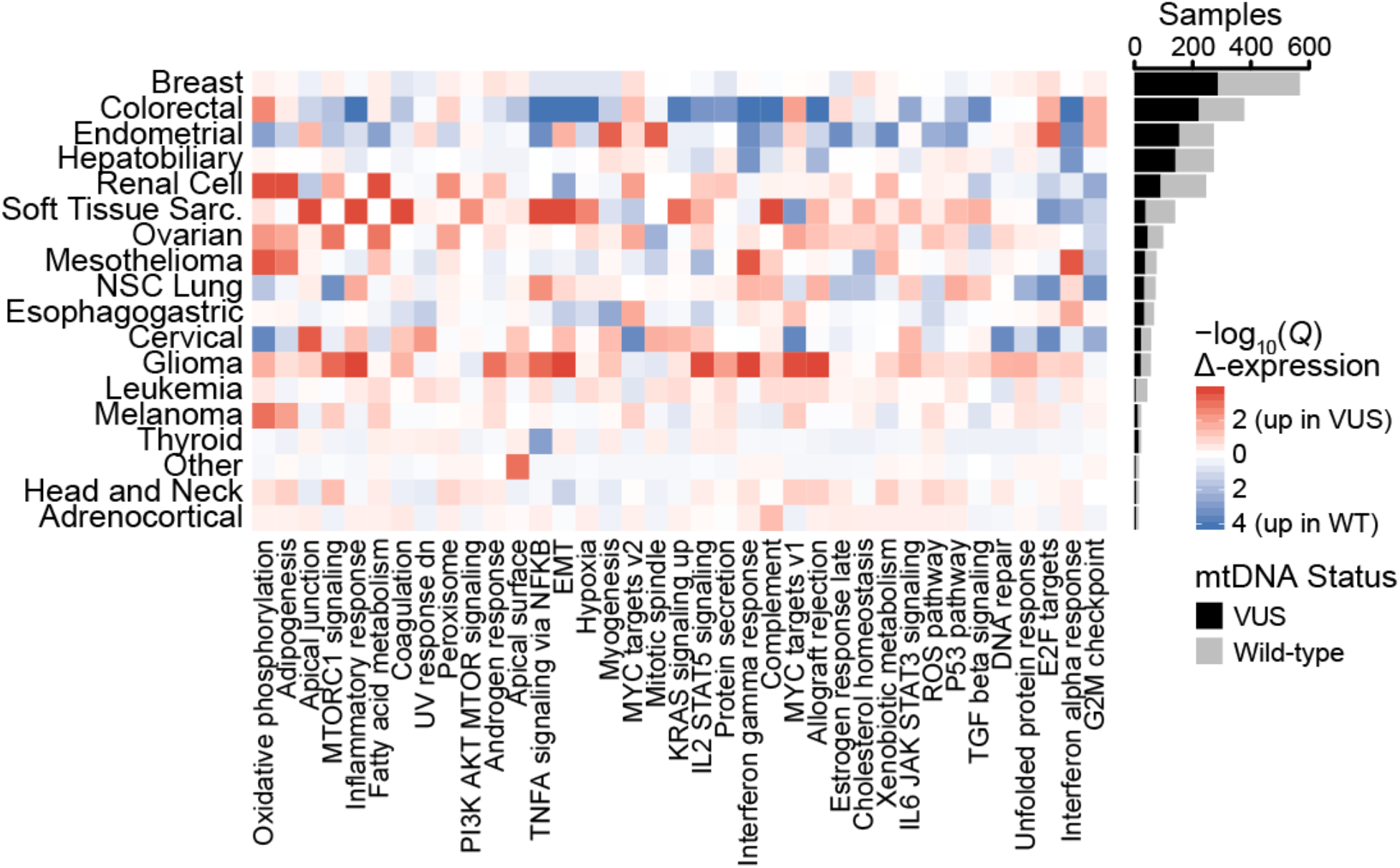
Transcriptional dysregulation attributed to mtDNA VUSs. Heatmap, differentially expressed mSigDB Hallmarks genesets between tumors with any somatic VUSs or wild-type mtDNA. Genesets ordered from most up-regulated across cancer types to most down-regulated. Barplot, number of cases with VUSs or wild-type mtDNA.

**Supplementary Fig. 4d:**
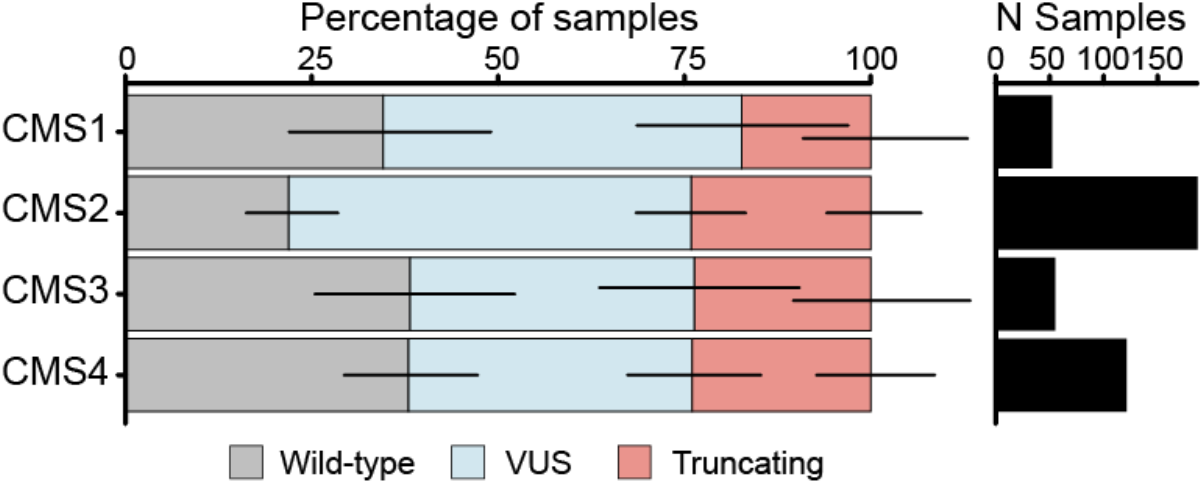
Difference in mtDNA mutation status between colorectal cancer consensus molecular subtypes. Left, the proportion of samples with wild-type mtDNA (*i.e.* no somatic mutations), VUS (any non-truncating) or truncating variants among colorectal tumors with each consensus molecular subtype (CMS) is shown. Right, histogram of the number of well-covered colorectal tumors. There was a statistically significant difference in mtDNA mutation status between different CMS classifications (*P*=0.03, Chi-squared test, *n*=415 samples total).

## Notes

### Competing Interest Statement

The authors have declared no competing interest.

